## Supplementary Information for "Inference of cell type-specific gene regulatory networks on cell lineages from single cell omic datasets"

Zhang et al.

### Supplementary Figures

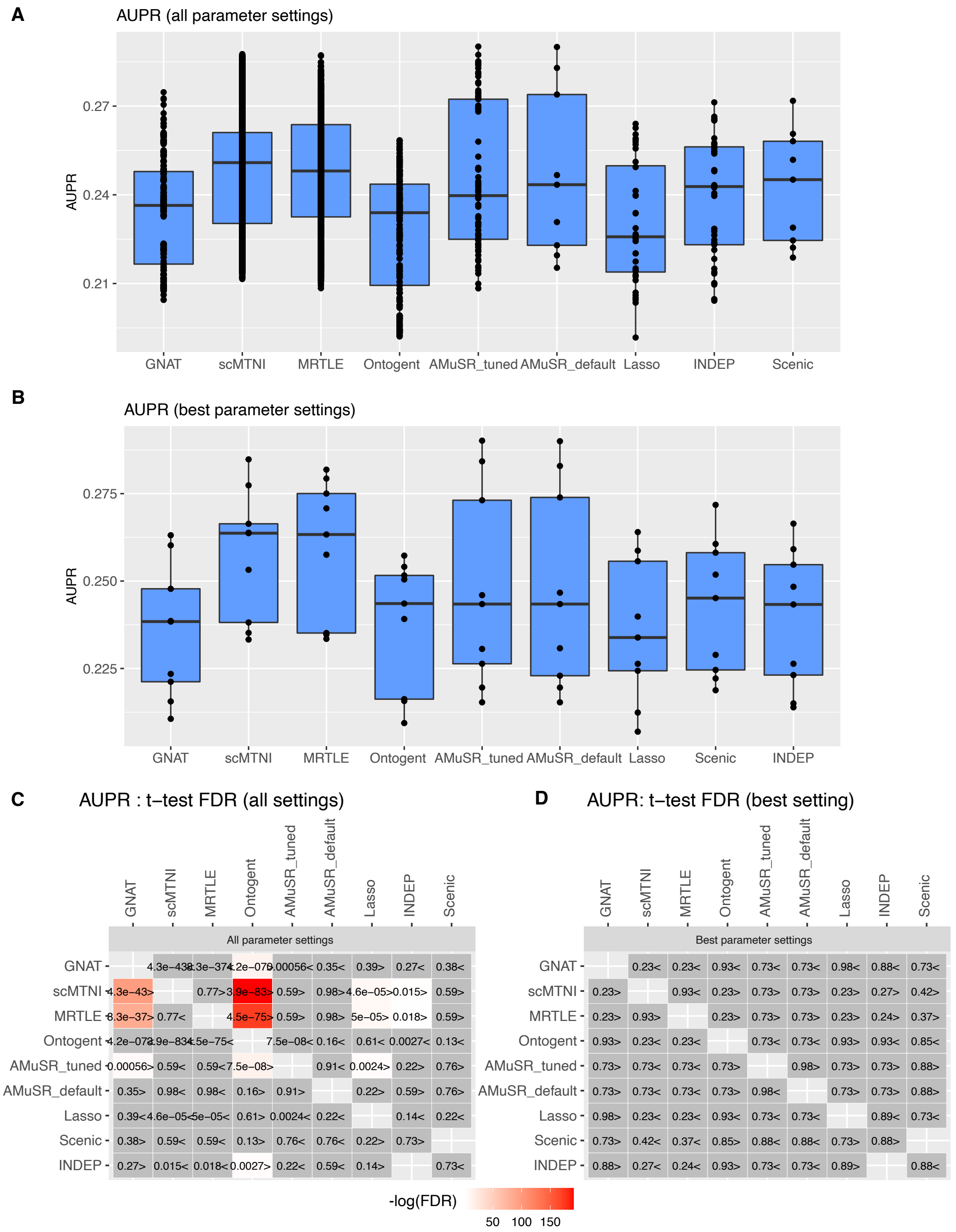

**Supplementary Figure 1.** Performance based on AUPR of different algorithms on three simulated datasets from three cell types. **A.** Boxplots show the AUPRs for all parameter settings of the tested algorithms. The number of experiments $n$ varies depending on the number of parameter settings per algorithm. **B.** Boxplots show the AUPRs for the best parameter setting, based on the AUPR, of the tested algorithms ($n = 9$, 3 cell types for 3 datasets). In both **A** and **B**, the horizontal middle line of each plot is the median. The bounds of the box are 0.25 quantile ($Q_1$) and 0.75 quantile ($Q_3$). The upper whisker is the minimum of the maximum value and $Q_3 + 1.5 * IQR$, where $IQR = Q_3 - Q_1$. The lower whisker is the maximum of the minimum value and $Q_1 - 1.5 * IQR$. **C.** T-test comparing the AUPR of the row algorithm to the column algorithm for all parameter settings. The two-sided t-test is conducted on AUPR of $n$ experiments for every pair of algorithms. For **A** and **C**, the total number of experiments for 3 cell types and 3 datasets is set as, GNAT:$n = 420$, scMTNI:$n = 11019$, MRTLE:$n = 1557$, Ontogent:$n = 528$, AMuSR_tuned:$n = 81$, AMuSR_default:$n = 9$, Lasso:$n = 36$, INDEP:$n = 45$, Scenic:$n = 9$. **D.** T-test comparing the AUPR of the row algorithm to the column algorithm for the best parameter setting. The two-sided t-test is conducted on AUPR of $n = 9$ experiments (3 cell types for 3 datasets) for every pair of algorithms. In both **C** and **D**, shown are FDR corrected P-values. The color scale is specified for $-log(FDR)$, with the red color proportional to significance.

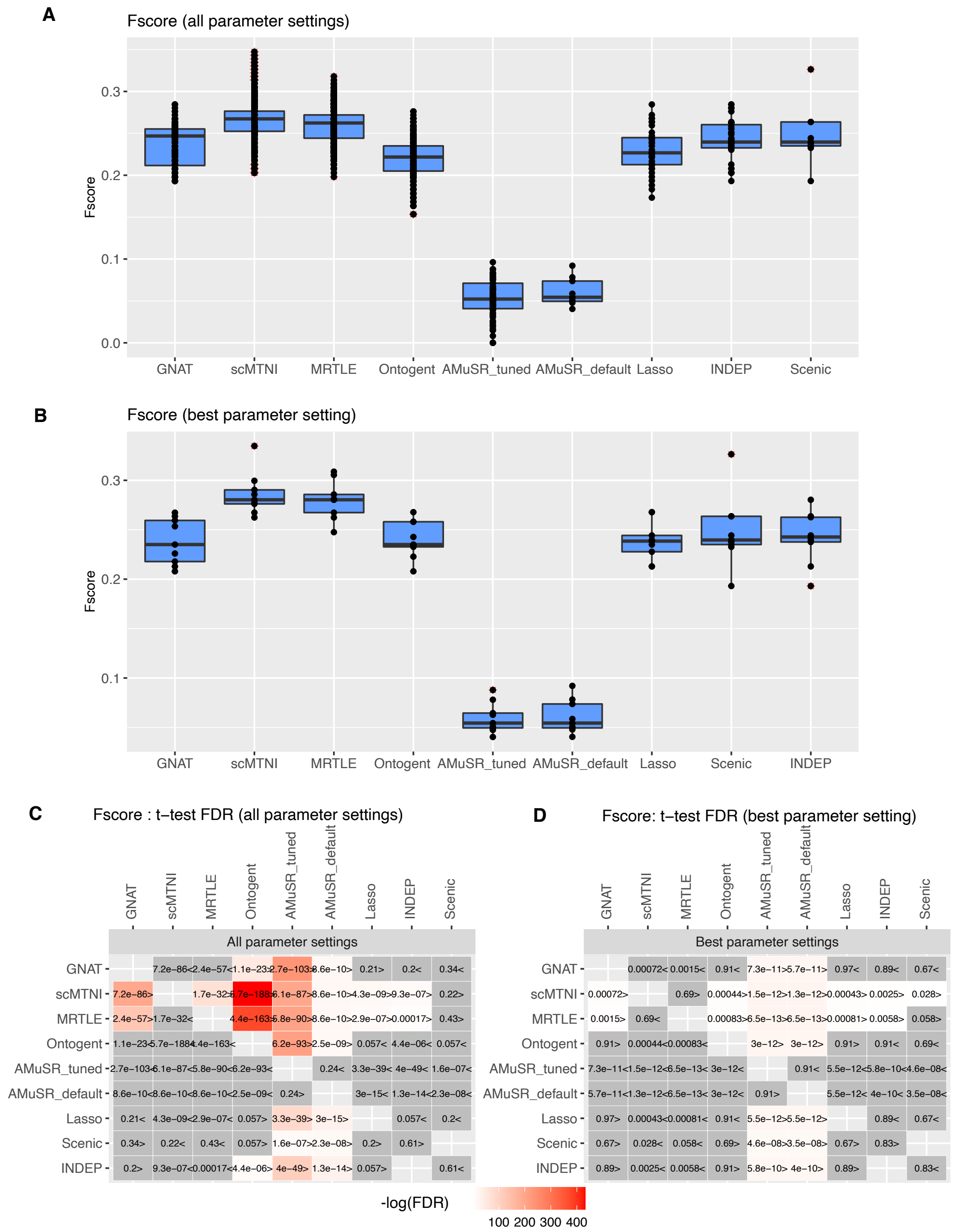

**Supplementary Figure 2.** Performance based on F-score of different algorithms on simulated data from three cell types. **A.** Boxplots show the F-score for all parameter settings of the tested algorithms. The number of experiments $n$ varies depending on the number of parameter settings per algorithm. **B.** Boxplots show the F-score for the best parameter setting, based on the Fscore, of the tested algorithms ($n = 9$, 3 cell types for 3 datasets). In both **A** and **B**, the horizontal middle line of each plot is the median. The bounds of the box are 0.25 quantile ($Q_1$) and 0.75 quantile ($Q_3$). The upper whisker is the minimum of the maximum value and $Q_3 + 1.5 * IQR$, where $IQR = Q_3 - Q_1$. The lower whisker is the maximum of the minimum value and $Q_1 - 1.5 * IQR$. **C.** T-test comparing the F-score of the row algorithm to the column algorithm for all parameter settings. The two-sided t-test is conducted on F-score of $n$ experiments for every pair of algorithms. For **A** and **C**, the total number of experiments for 3 cell types and 3 datasets is set as, GNAT:$n = 420$, scMTNI:$n = 11019$, MRTLE:$n = 1557$, Ontogent:$n = 528$, AMuSR_tuned:$n = 81$, AMuSR_default:$n = 9$, Lasso:$n = 36$, INDEP:$n = 45$, Scenic:$n = 9$. **D.** T-test comparing the F-score of the row algorithm to the column algorithm for the best parameter setting. The two-sided t-test is conducted on F-score of $n = 9$ experiments (3 cell types for 3 datasets) for every pair of algorithms. In both C and D, shown are FDR corrected P-values. The color scale is specified for $-log(FDR)$, with the red color proportional to significance.

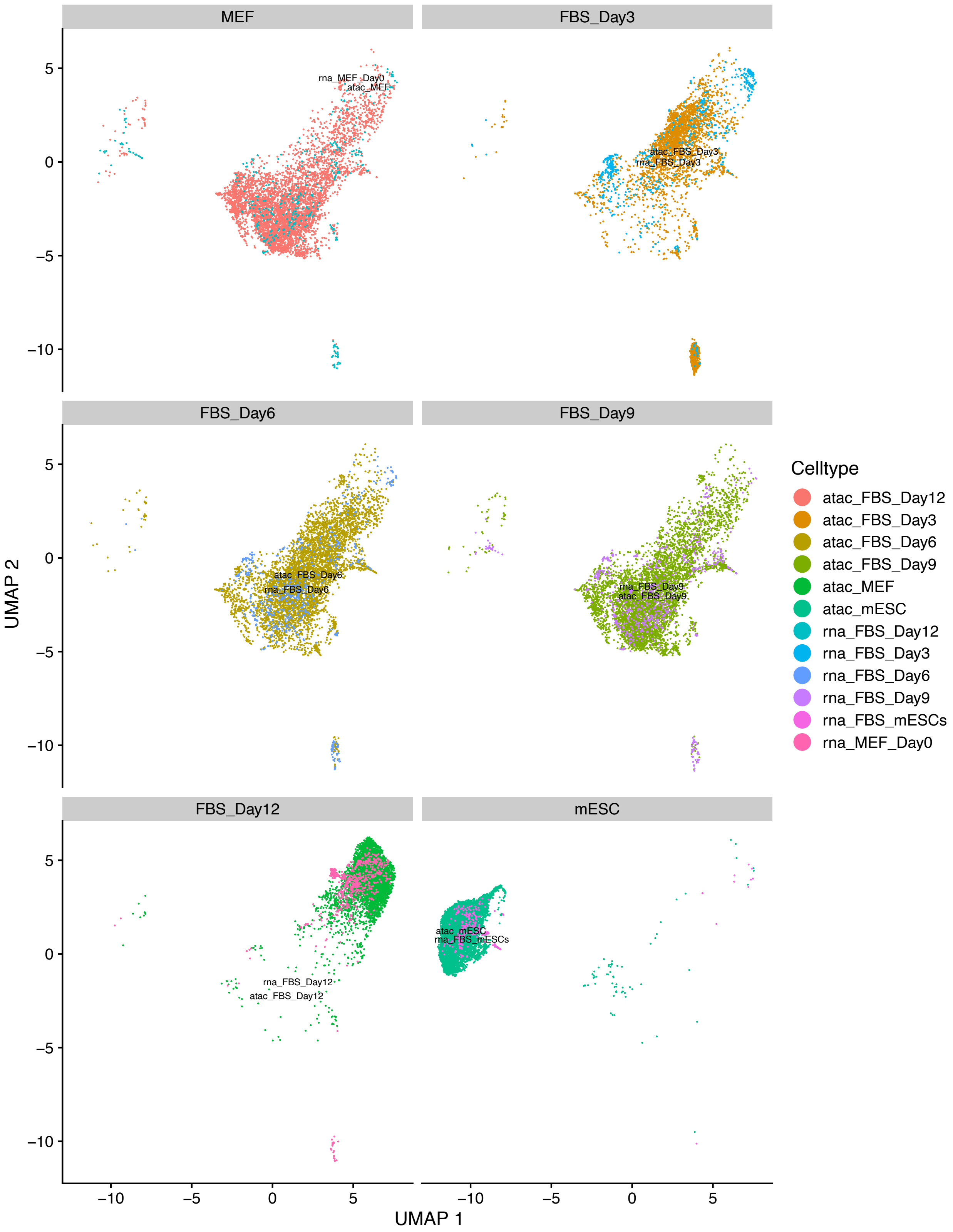

**Supplementary Figure 3.** UMAP depicting the sample labels of the scATAC-seq and scRNA-seq data separated by each sample for the FBS reprogramming data.

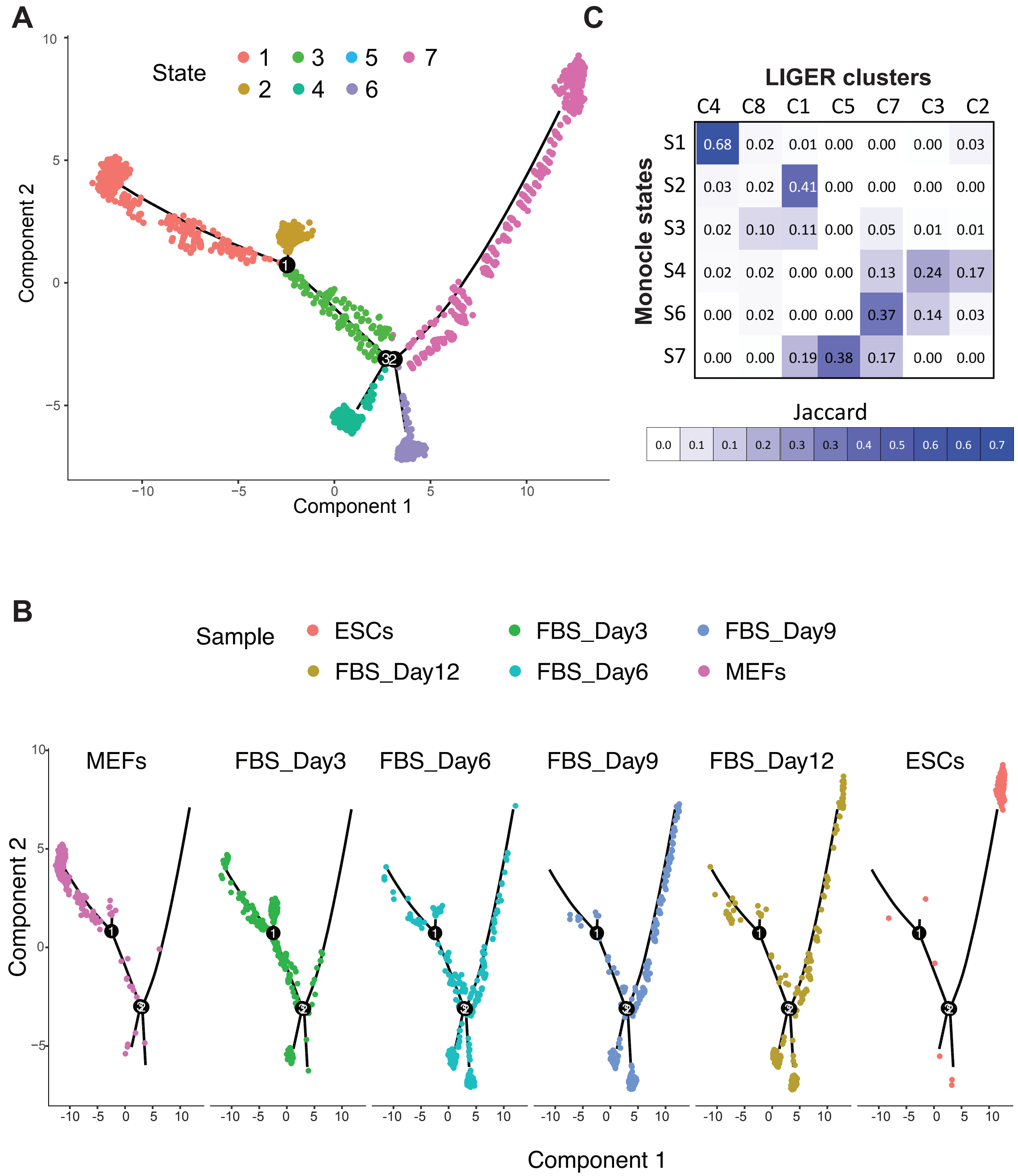

**Supplementary Figure 4.** Monocle trajectory analysis of FBS reprogramming scRNA-seq data. **A.** Monocle state diagram exhibiting the trajectory of cells from a differentiated state (State 1, MEFs), to a pluripotent state (State 7, ESCs). States 4 and 6 represent cells that do not reach the pluripotent end point. **B.** Overlay of cells from individual samples onto the Monocle trajectory. **C.** Overlap as measured by Jaccard coefficient between Monocle states (row) and LIGER cell clusters (columns). Each entry shows the Jaccard coefficient score for the overlap of a state and cell cluster. The higher the blue intensity the greater the overlap.

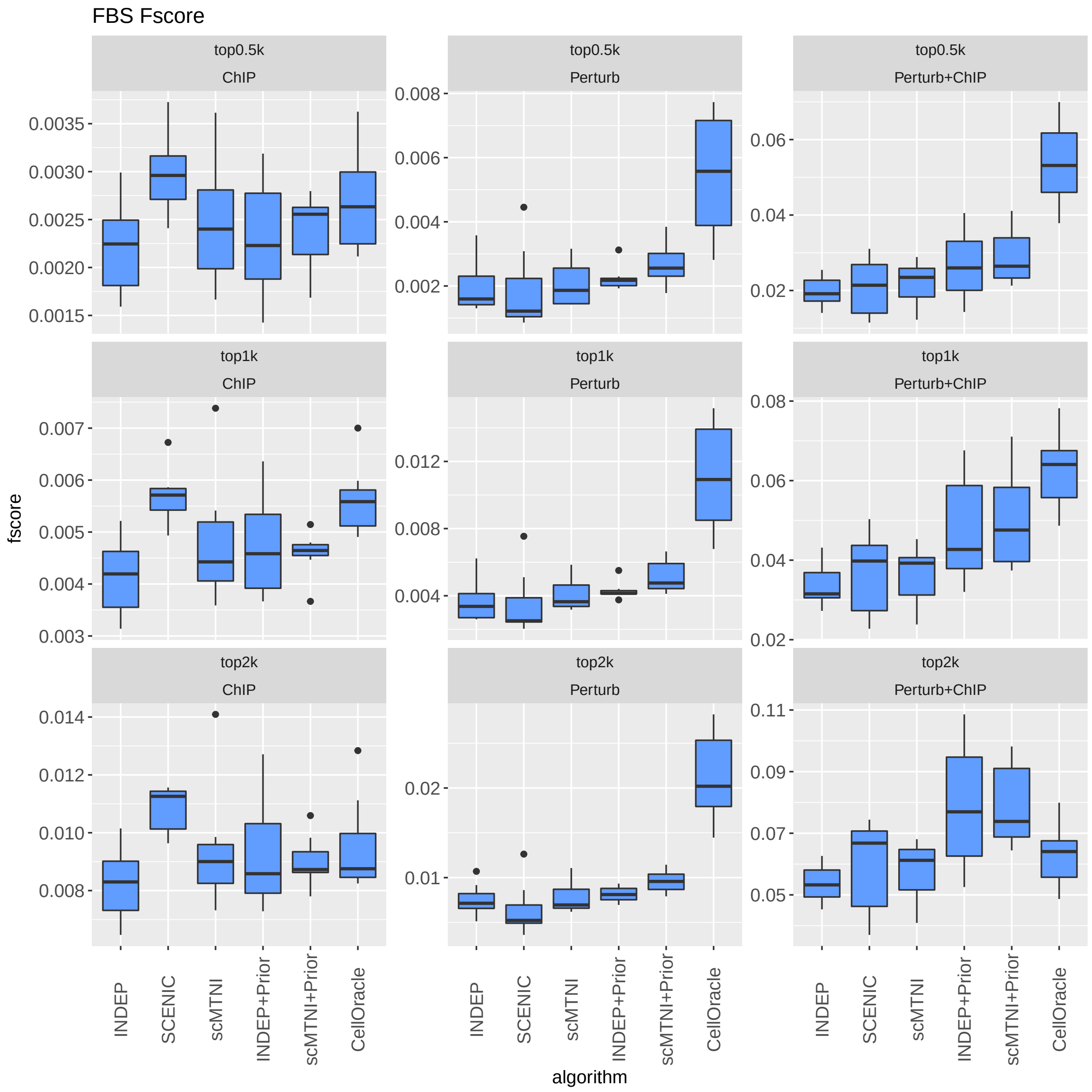

**Supplementary Figure 5.** F-score of $n = 7$ cell clusters for top 500, 1k, 2k edges in predicted networks of scMTNI, scMTNI with prior (scMTNI+Prior), INDEP, INDEP with prior (INDEP+Prior), SCENIC and CellOracle on mouse cellular reprogramming data compared to three gold standard datasets: ChIP, Perturb and Perturb+ChIP. In the boxplot, the horizontal middle line of each plot is the median. The bounds of the box are 0.25 quantile ($Q_1$) and 0.75 quantile ($Q_3$). The upper whisker is the minimum of the maximum value and $Q_3 + 1.5 * IQR$, where $IQR = Q_3 - Q_1$. The lower whisker is the maximum of the minimum value and $Q_1 - 1.5 * IQR$.

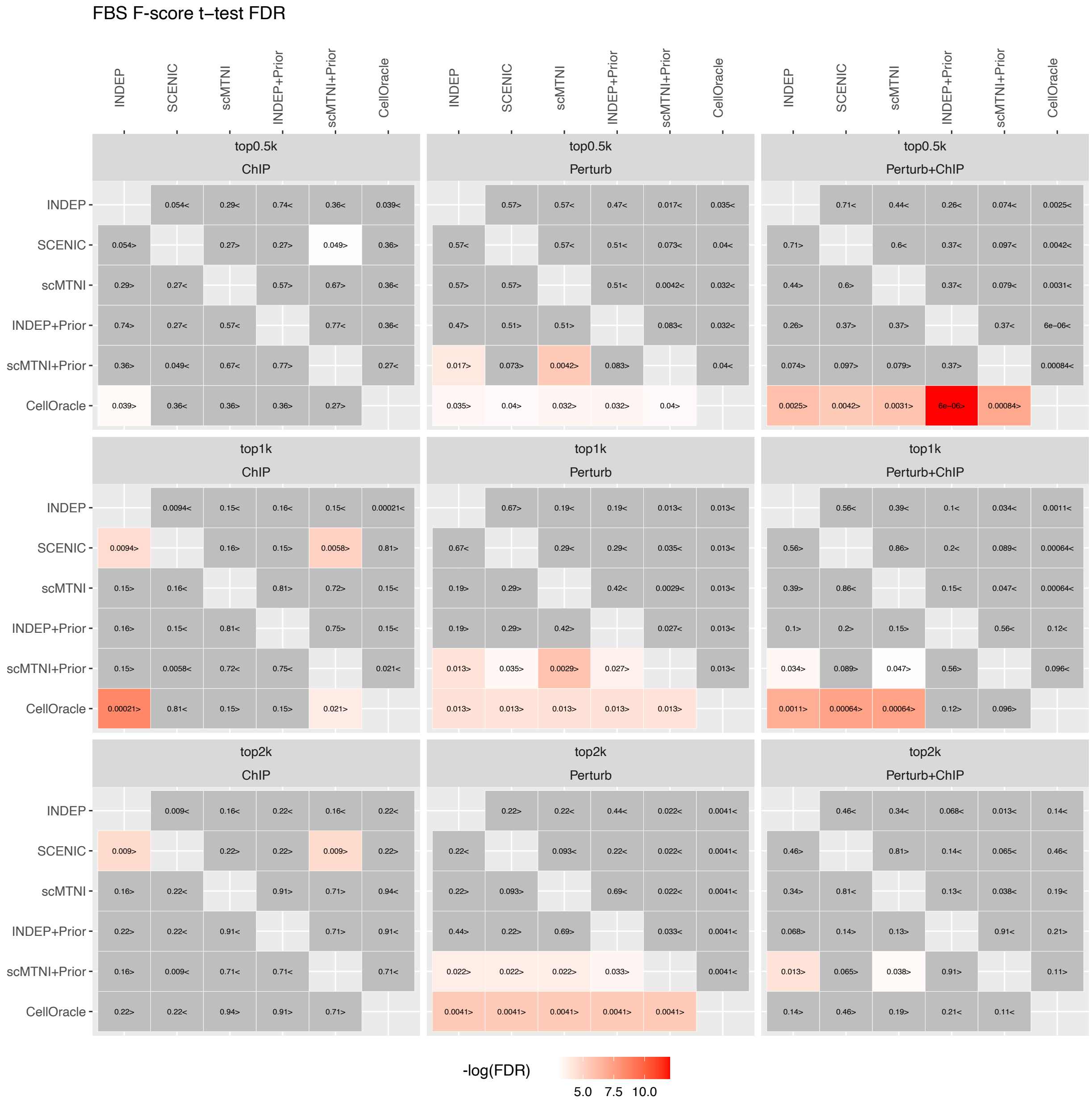

**Supplementary Figure 6.** Heatmap of FDR corrected T-test p-values comparing the F-score of top 500, 1k, 2k edges in predicted networks of scMTNI, scMTNI+Prior, INDEP, INDEP+Prior, SCENIC and CellOracle on mouse reprogramming data compared to the three gold standard datasets. The two-sided paired t-test is conducted on F-scores of $n = 7$ cell clusters for every pair of algorithms, comparing whether the row algorithm's F-score is higher than the column algorithm's F-score. Significant difference (FDR$<$0.05) is highlighted using a white-red colormap (the color scale for $-\log(\text{FDR})$). Non-significance is colored in gray. The sign"$<$" or "$>$" specifies whether the row algorithm's F-scores were worse or better than the column algorithm's F-scores.

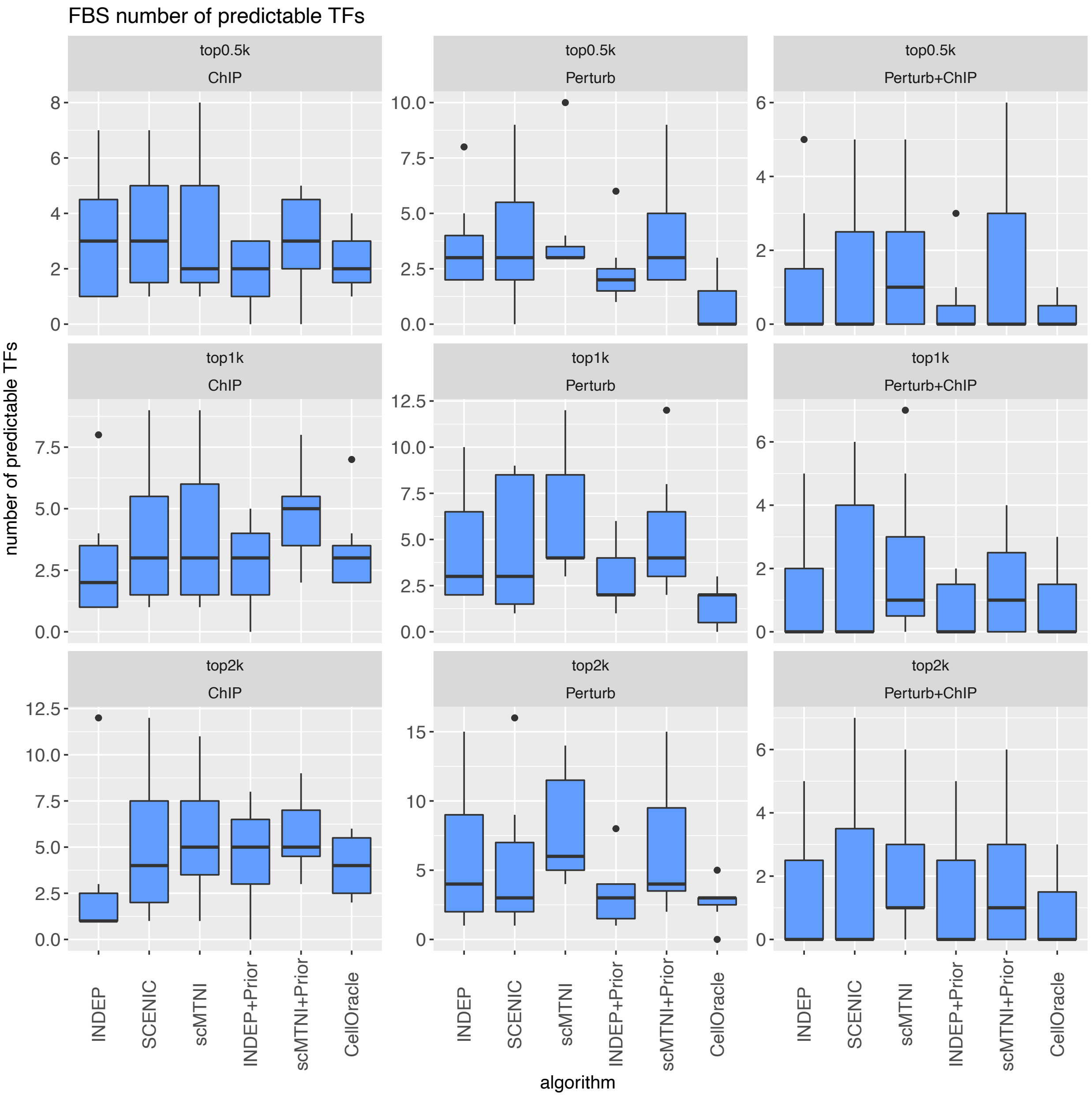

**Supplementary Figure 7.** Number of predictable TFs in top 500, 1k, 2k edges in predicted networks of scMTNI, scMTNI with prior (scMTNI+Prior), INDEP, INDEP with prior (INDEP+Prior), SCENIC and CellOracle on the mouse reprogramming data. Shown are the predictable TF numbers of $n = 7$ cell clusters for the three gold standard datasets, ChIP, Perturb and Pertrub+ChIP. In the boxplot, the horizontal middle line of each plot is the median. The bounds of the box are 0.25 quantile ($Q_1$) and 0.75 quantile ($Q_3$). The upper whisker is the minimum of the maximum value and $Q_3 + 1.5 * IQR$, where $IQR = Q_3 - Q_1$. The lower whisker is the maximum of the minimum value and $Q_1 - 1.5 * IQR$.

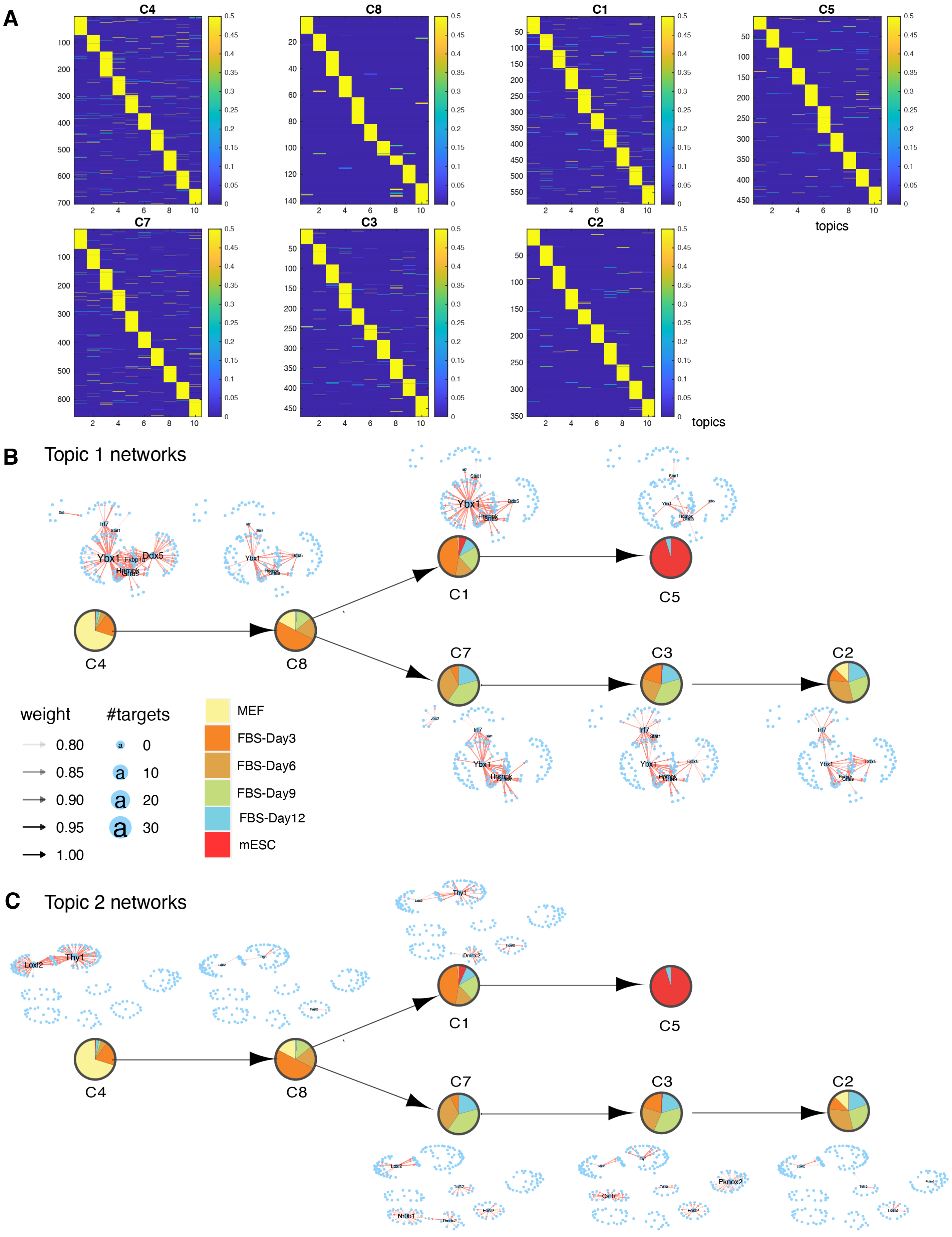

**Supplementary Figure 8.** LDA analysis of scMTNI+Prior inferred networks on mouse reprogramming data. **A.** The document-topic weight matrix for all genes for all cell clusters with 10 topics. Rows correspond to regulators (documents) and columns correspond to topics. Rows are ordered based on topic membership. **B.** Topic-specific networks for each cell cluster for topic 1. **C.** Topic-specific networks for each cell cluster for topic 2.

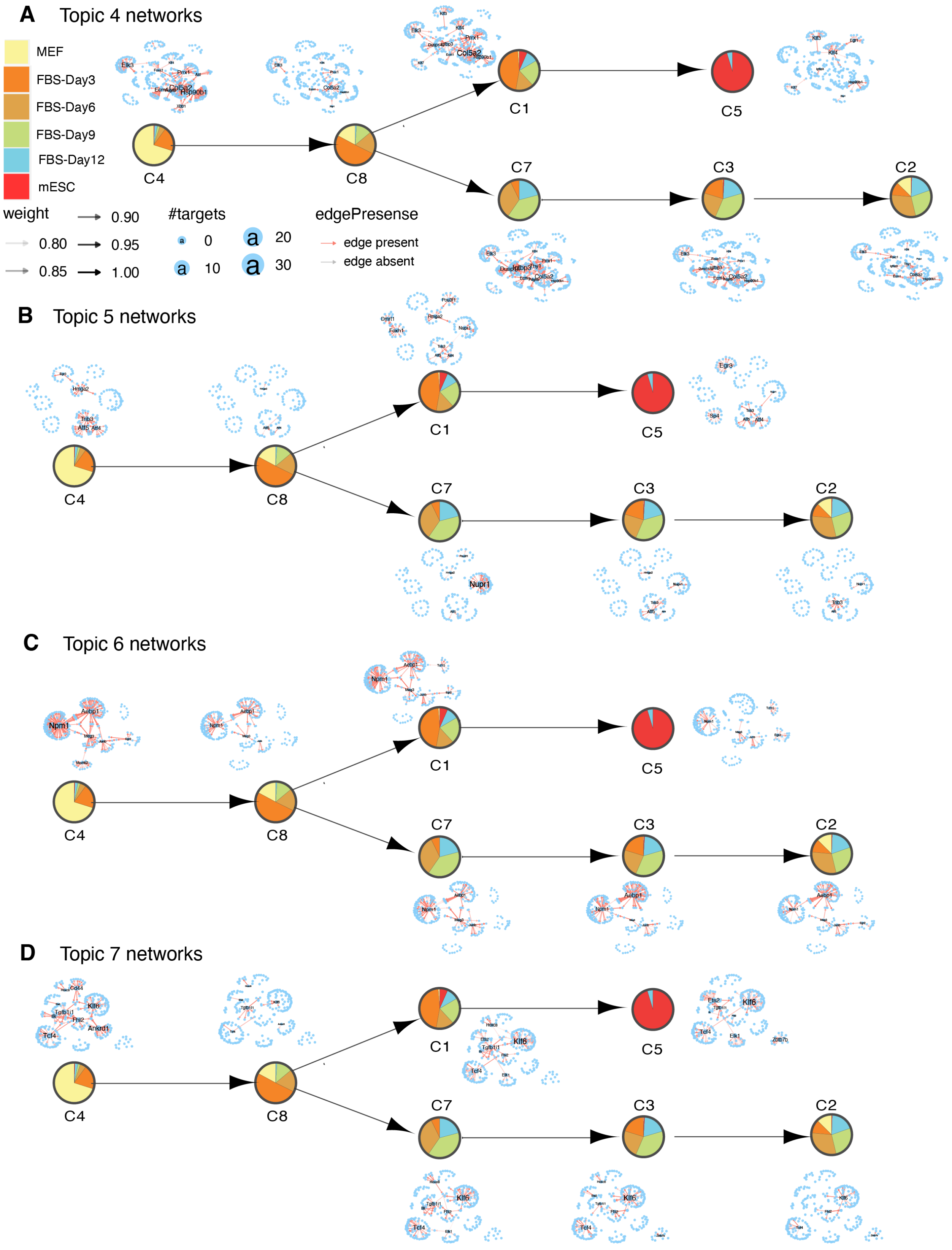

**Supplementary Figure 9.** Topic-specific networks for each cell cluster identified from scMTNI networks on mouse reprogramming data for **A.** topic 4, **B.** topic 5, **C.** topic 6, and **D** topic 7.

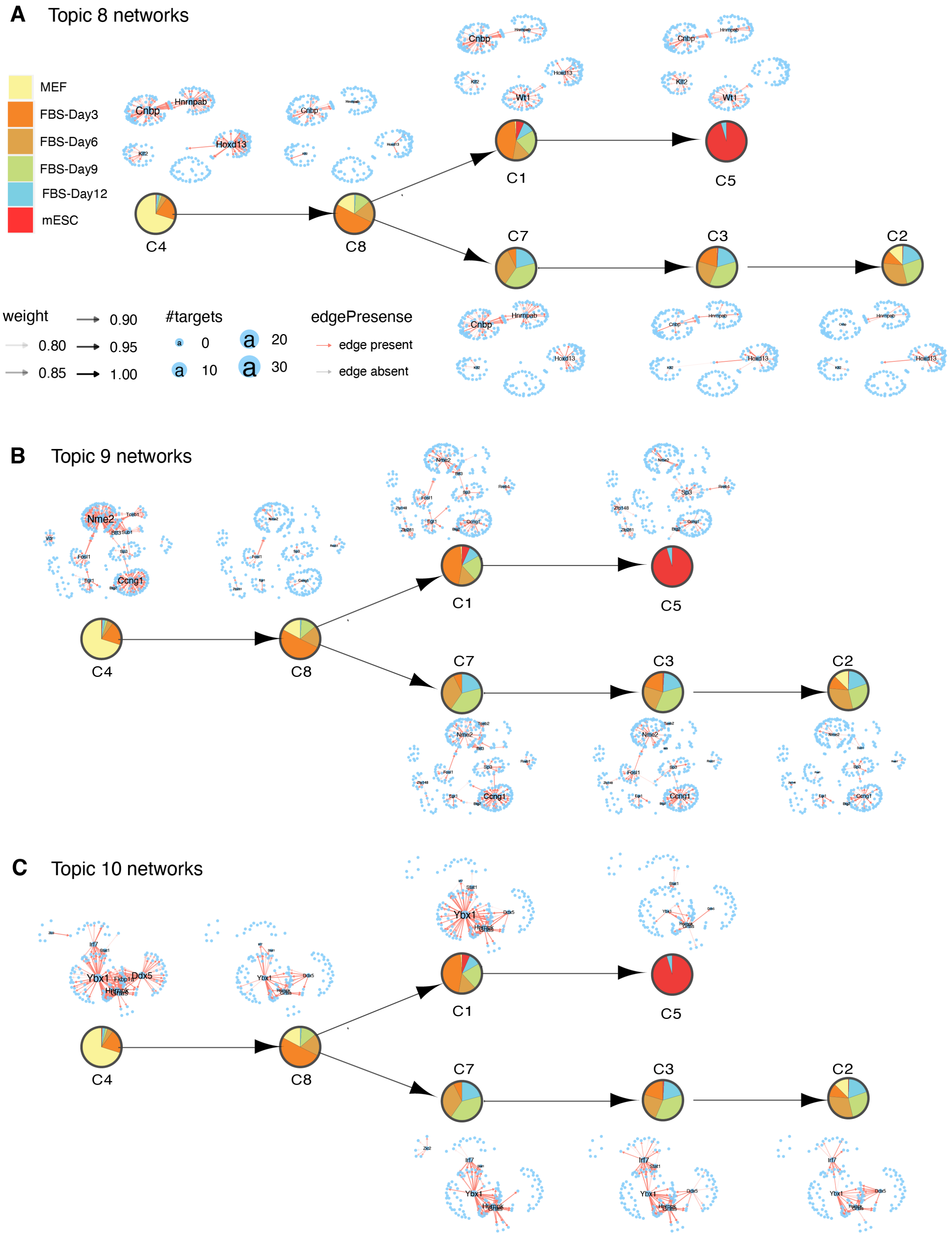

**Supplementary Figure 10.** Topic-specific networks for each cell cluster identified from scMTNI networks on mouse reprogramming data for **A.** topic 8, **B.** topic 9, and **C.** topic 10.

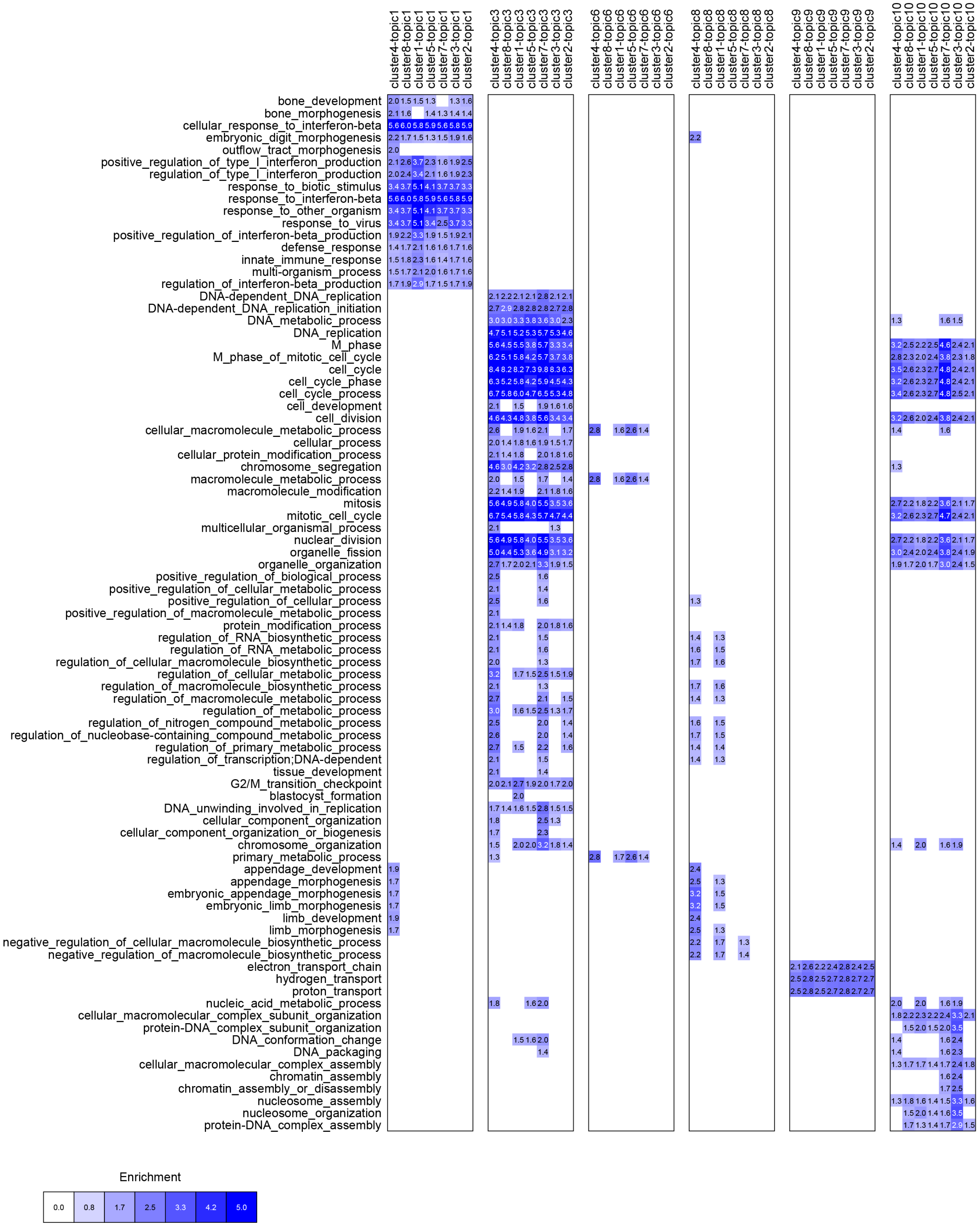

**Supplementary Figure 11.** Gene Ontology (GO) process enrichment for genes associated with each LDA topic-specific network identified from mouse reprogramming data. We used FDR<0.01 to determine significantly enriched terms. The blue intensity of the heatmap corresponds to $-\log10(\text{FDR})$.

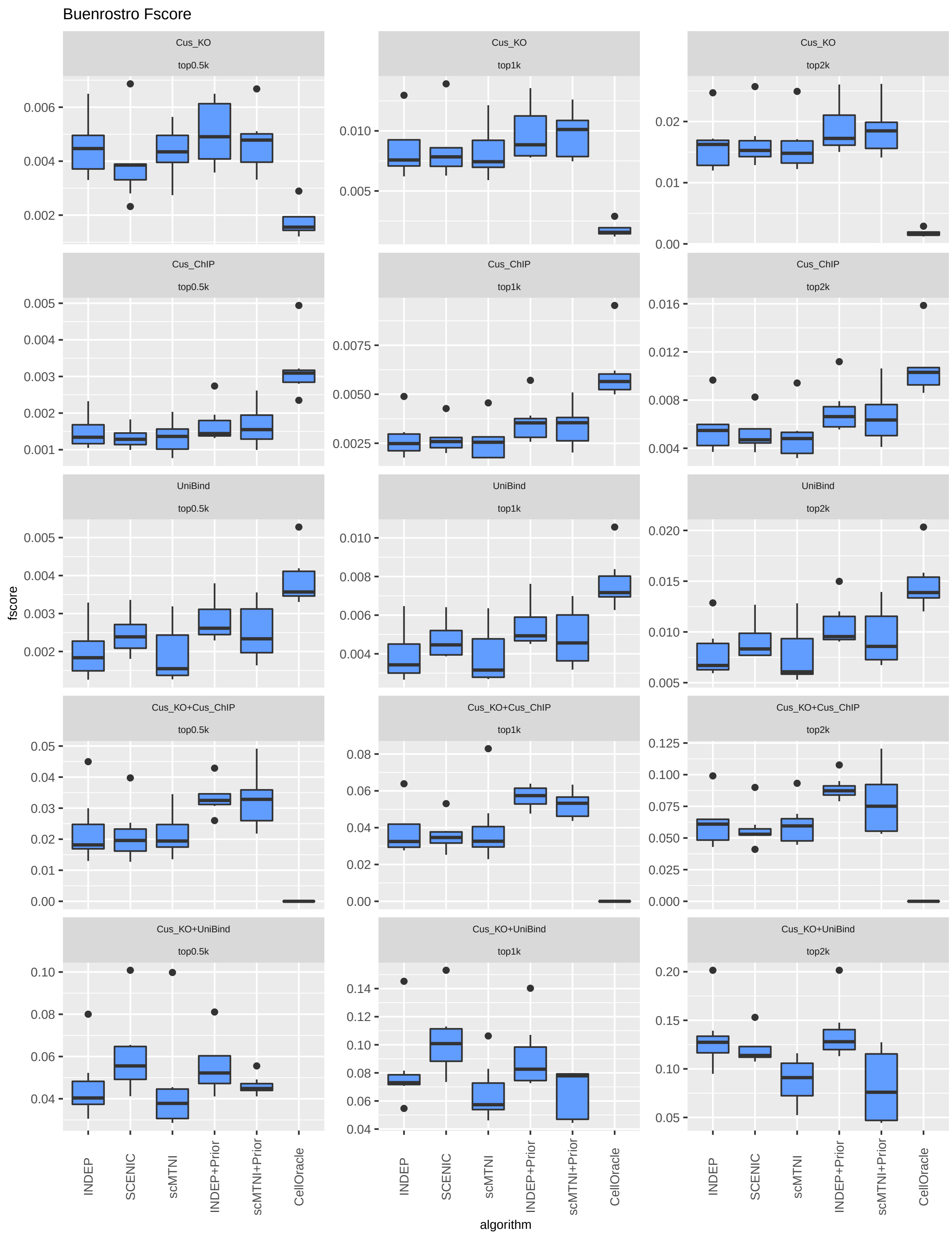

**Supplementary Figure 12.** F-score of top 500, 1k, 2k edges in predicted networks of scMTNI, scMTNI+Prior, INDEP, INDEP+Prior, SCENIC and CellOracle on human hematopoietic differentiation data from Buenrostro et al. Shown are results of $n = 7$ cell clusters (all cell clusters excluding C1) for five different gold standards (rows). In the boxplot, the horizontal middle line of each plot is the median. The bounds of the box are 0.25 quantile ($Q_1$) and 0.75 quantile ($Q_3$). The upper whisker is the minimum of the maximum value and $Q_3 + 1.5 * IQR$, where $IQR = Q_3 - Q_1$. The lower whisker is the maximum of the minimum value and $Q_1 - 1.5 * IQR$.

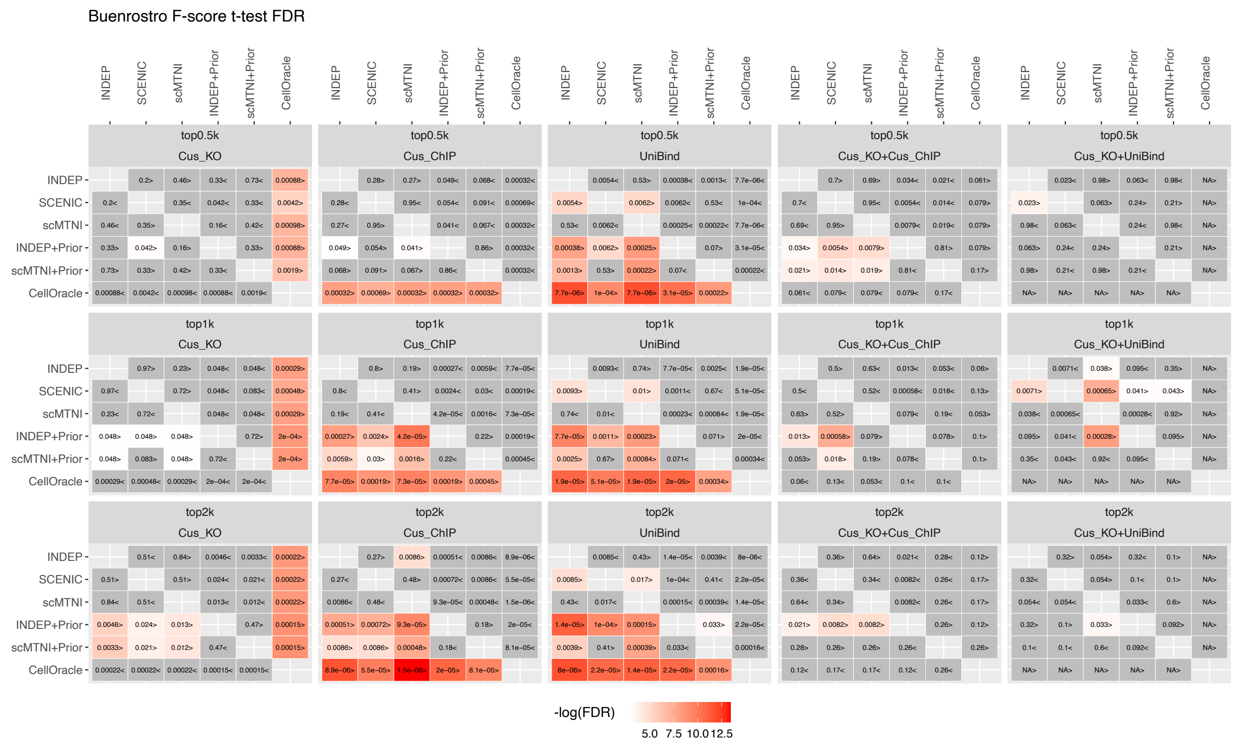

**Supplementary Figure 13.** Heatmap of FDR corrected T-test pvalues comparing the F-score of top 500, 1k, 2k edges in predicted networks of scMTNI, scMTNI+Prior, INDEP, INDEP+Prior, SCENIC and CellOracle on human hematopoietic differentiation data from Buenrostro et al. Shown are relative performance using five gold standard datasets. The two-sided paired t-test is conducted on F-scores of $n = 7$ cell clusters (all cell clusters excluding C1) for every pair of algorithms, comparing whether the row algorithm's F-score is higher than the column algorithm's F-score. Significant difference (FDR<0.05) is highlighted using a white-red colormap (the color scale for $-\log(\text{FDR})$). Non-significance is colored in gray. The sign"<" or ">" specifies whether the row algorithm's F-scores were worse or better than the column algorithm's F-scores.

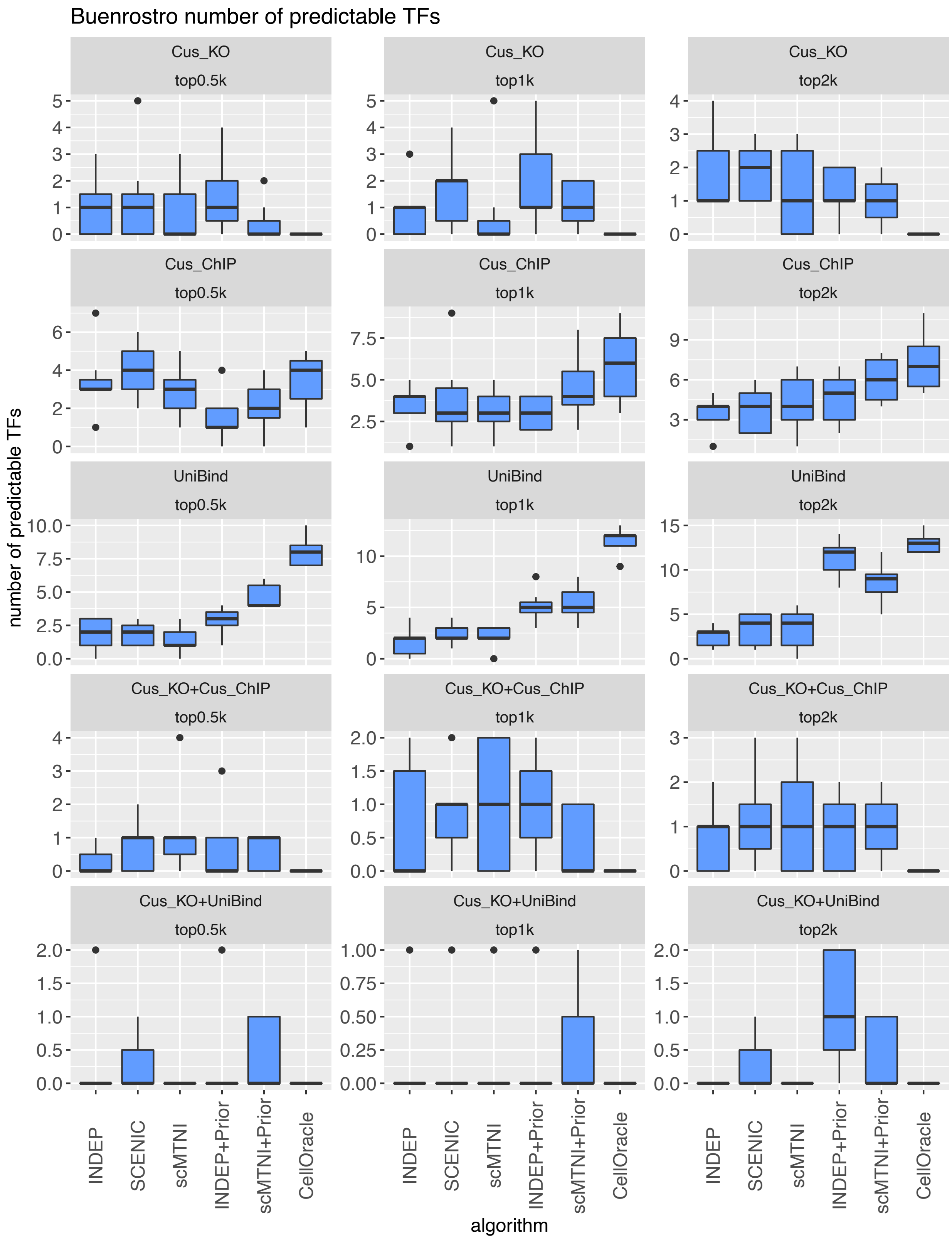

**Supplementary Figure 14.** Number of predictable TFs in top 500, 1k, 2k edges in predicted networks of scMTNI, scMTNI+Prior, INDEP, INDEP+Prior, SCENIC and CellOracle on human hematopoietic differentiation data. Predictable TFs of $n = 7$ cell clusters (all cell clusters excluding C1) are computed on five gold standard datasets. In the boxplot, the horizontal middle line of each plot is the median. The bounds of the box are 0.25 quantile ($Q_1$) and 0.75 quantile ($Q_3$). The upper whisker is the minimum of the maximum value and $Q_3 + 1.5 * IQR$, where $IQR = Q_3 - Q_1$. The lower whisker is the maximum of the minimum value and $Q_1 - 1.5 * IQR$.

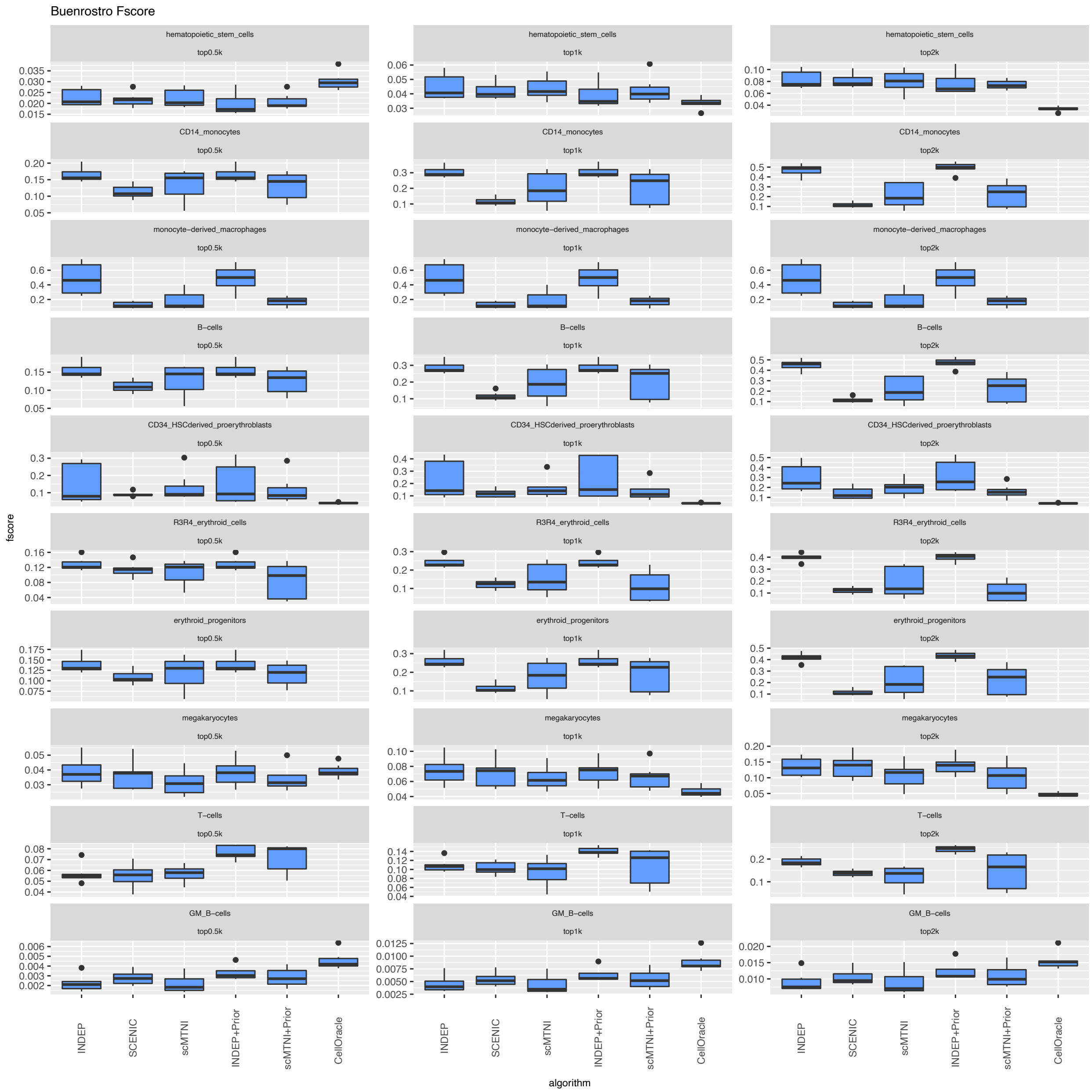

**Supplementary Figure 15.** F-score of $n = 7$ cell clusters (all cell clusters excluding C1) for top 500, 1k, 2k edges in predicted networks of scMTNI, scMTNI+Prior, INDEP, INDEP+Prior, SCENIC and CellOracle on human hematopoietic differentiation data for individual cell type-specific gold standard datasets. Each row corresponds to a particular gold standard and column corresponds to the number of top edges. In the boxplot, the horizontal middle line of each plot is the median. The bounds of the box are 0.25 quantile ($Q_1$) and 0.75 quantile ($Q_3$). The upper whisker is the minimum of the maximum value and $Q_3 + 1.5 * IQR$, where $IQR = Q_3 - Q_1$. The lower whisker is the maximum of the minimum value and $Q_1 - 1.5 * IQR$.

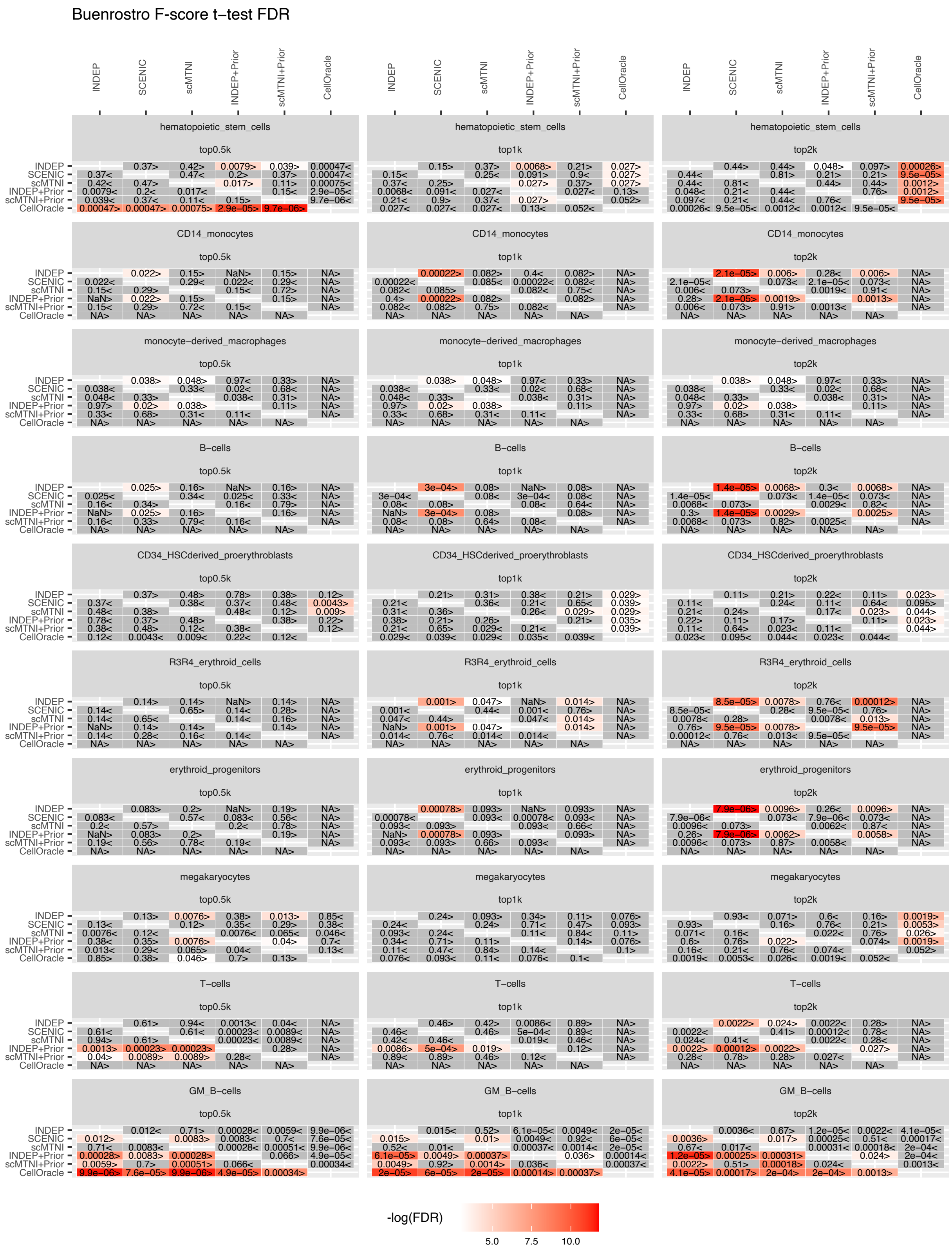

**Supplementary Figure 16.** Heatmap of FDR corrected t-test P-value to compare the F-score of top 500, 1k, 2k edges in predicted networks of scMTNI, scMTNI+Prior, INDEP, INDEP+Prior, SCENIC and CellOracle on human hematopoietic differentiation data. F-scores are computed on cell type specific gold standard datasets. The two-sided paired t-test is conducted on F-scores of $n = 7$ cell clusters (all cell clusters excluding C1) for every pair of algorithms, comparing whether the row algorithm's F-score is higher than the column algorithm's F-score. Significant difference (FDR$<$0.05) is highlighted using a white-red colormap (the color scale for $-\log(\text{FDR})$). Non-significance is colored in gray. The sign"$<$" or "$>$" specifies whether the row algorithm's F-scores were worse or better than the column algorithm's F-scores.

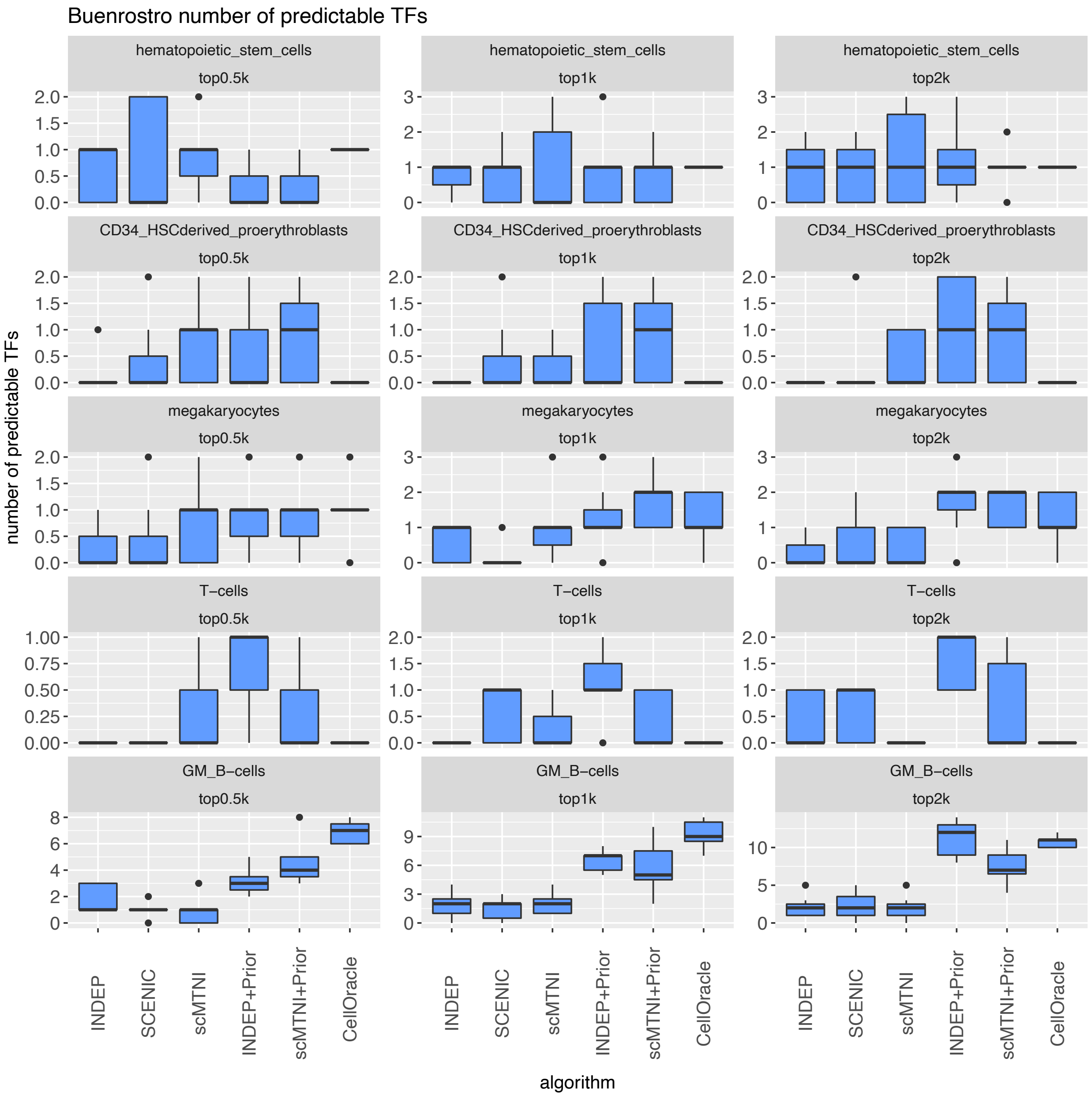

**Supplementary Figure 17.** Number of predictable TFs in top 500, 1k, 2k edges in predicted networks of scMTNI, scMTNI+Prior, INDEP, INDEP+Prior, SCENIC and CellOracle on human hematopoietic differentiation data. Predictable TFs of $n = 7$ cell clusters (all cell clusters excluding C1) are estimated for individual cell type-specific gold standard datasets. Each row corresponds to a particular gold standard and column corresponds to the number of top edges. Only gold standards that have any predictable TFs are shown. In the boxplot, the horizontal middle line of each plot is the median. The bounds of the box are 0.25 quantile ($Q_1$) and 0.75 quantile ($Q_3$). The upper whisker is the minimum of the maximum value and $Q_3 + 1.5 * IQR$, where $IQR = Q_3 - Q_1$. The lower whisker is the maximum of the minimum value and $Q_1 - 1.5 * IQR$.

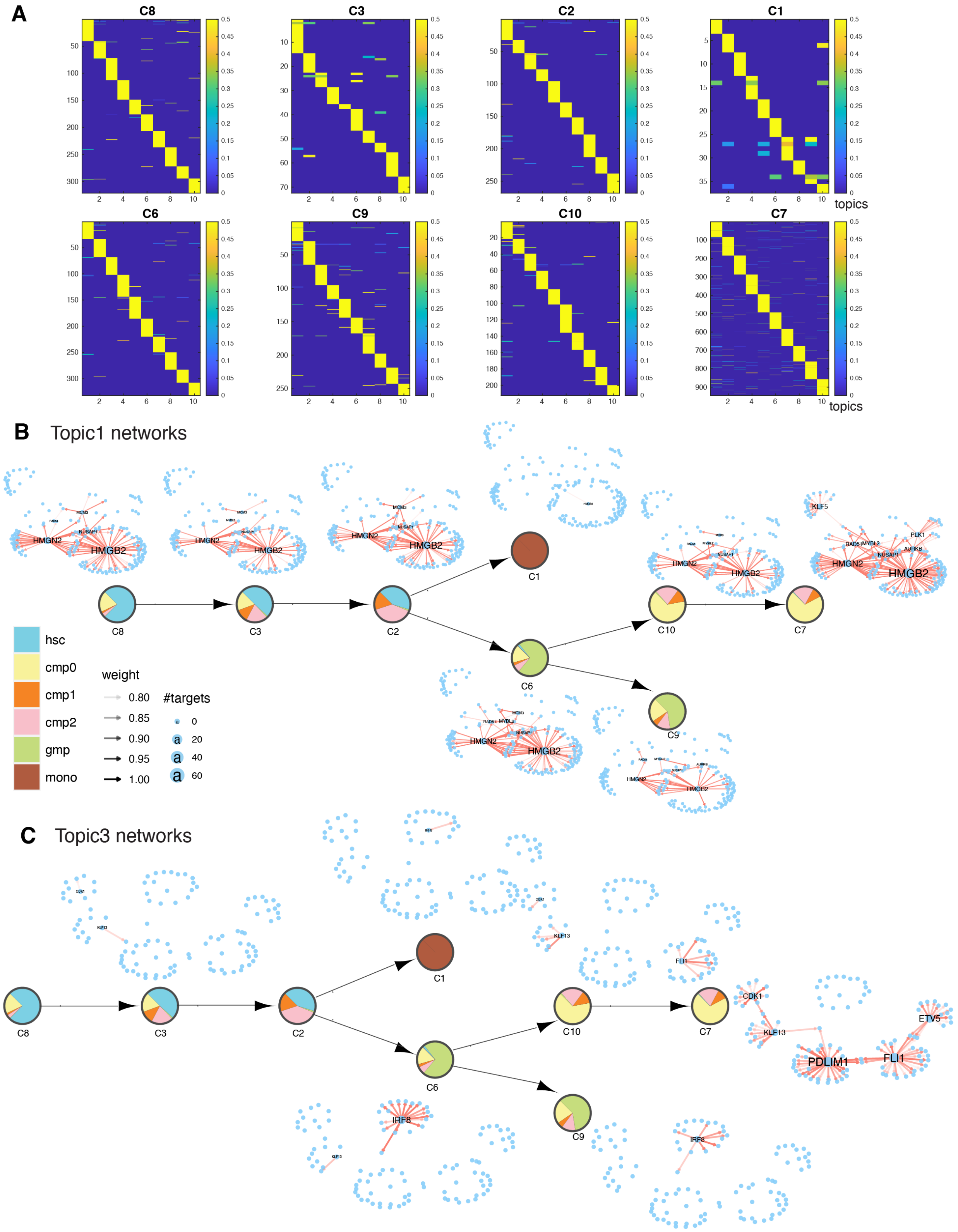

**Supplementary Figure 18.** LDA analysis of scMTNI inferred networks on human hematopoietic differentiation data from Buenrostro et al. **A.** The document-topic weight matrix for all regulators for all cell clusters with 10 topics. Topic-specific networks for each cell cluster for topic 1 (**B**) and topic 3 (**C**).

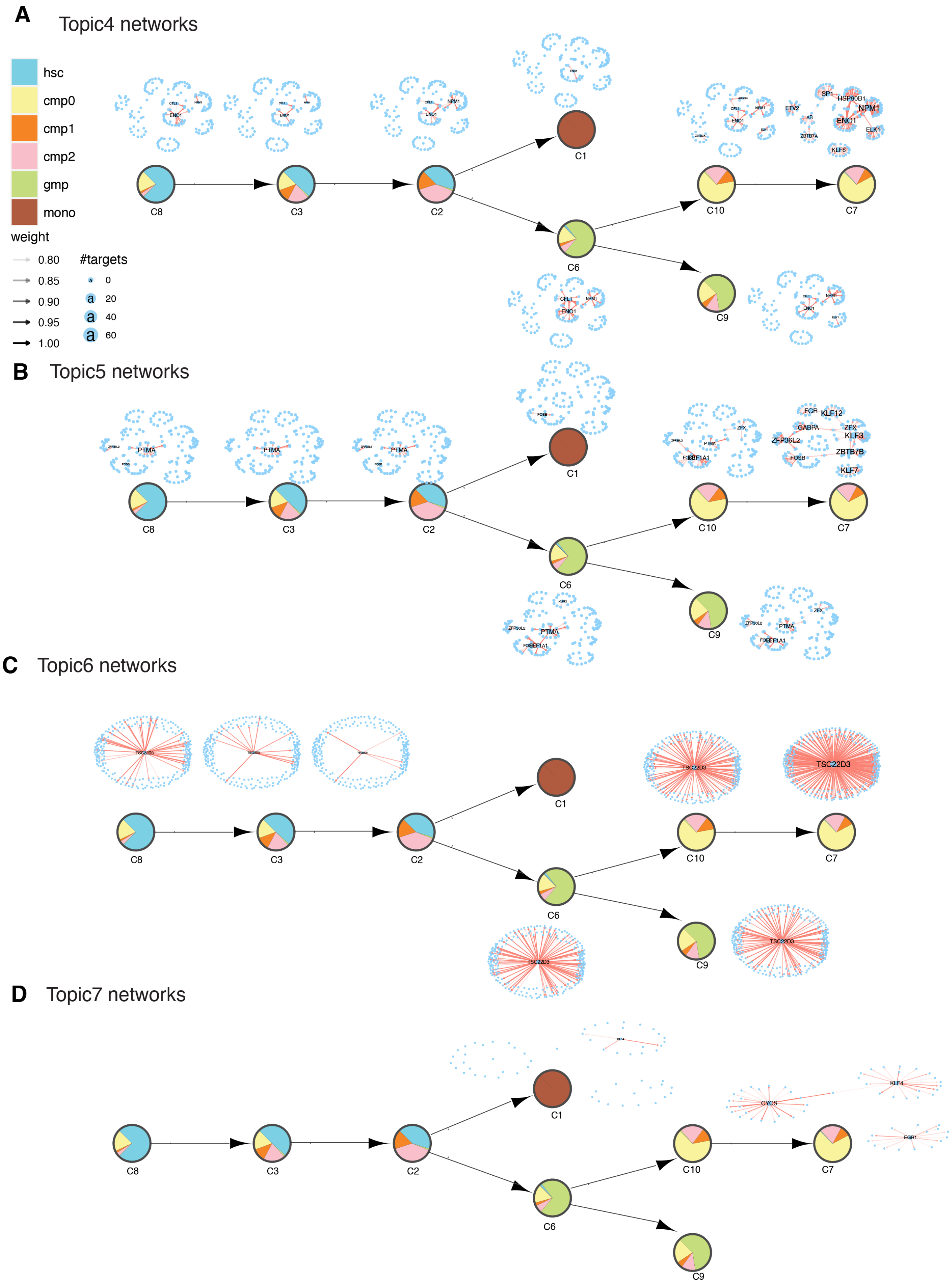

**Supplementary Figure 19.** Topic-specific networks obtained from scMTNI inferred networks on human hematopoietic differentiation data from Buenrostro et al. for topic 4 (**A**), topic 5 (**B**), topic 6 (**C**) and topic 7 (**D**).

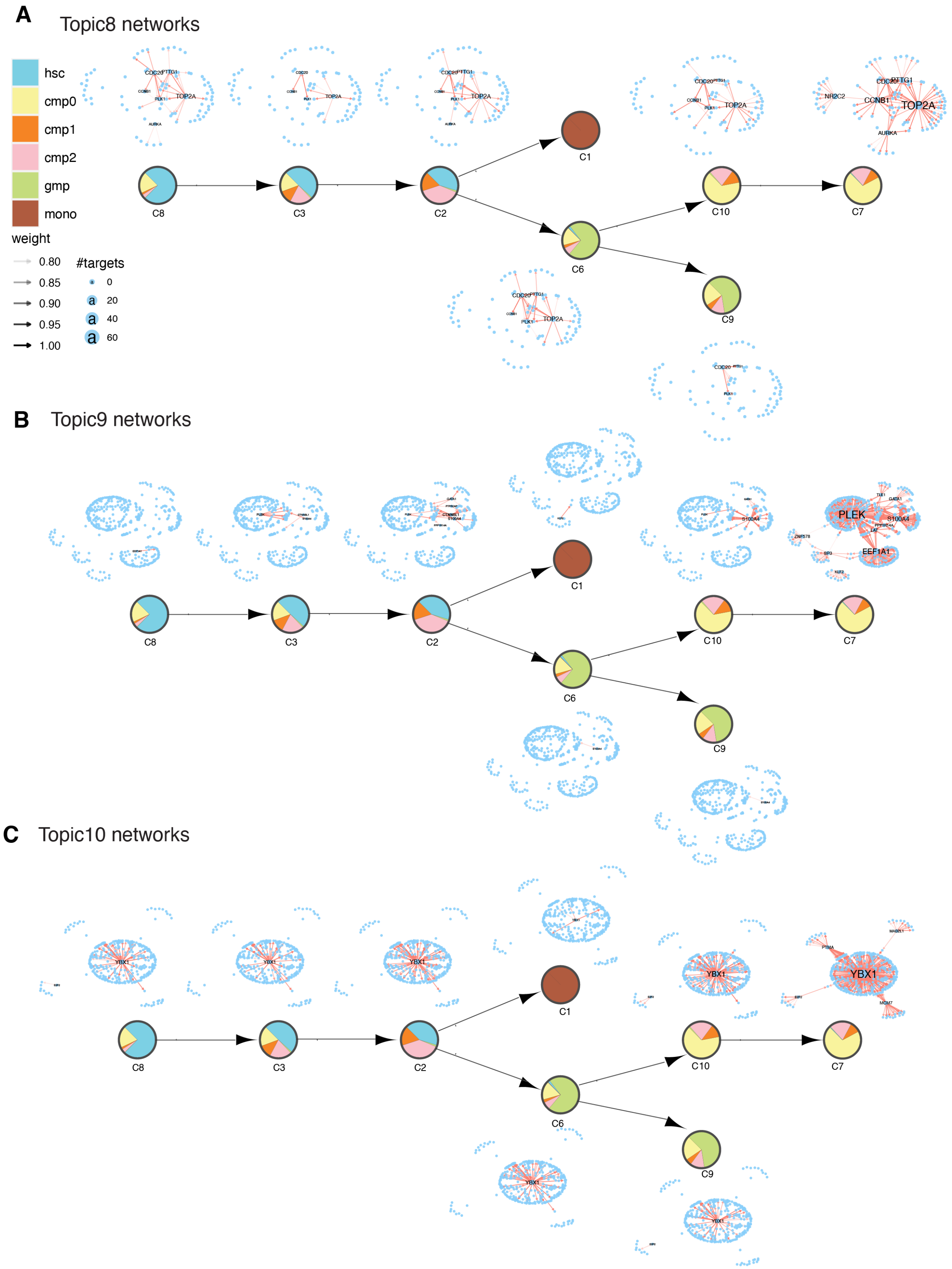

**Supplementary Figure 20.** Topic-specific networks from scMTNI inferred networks on human hematopoietic differentiation data from Buenrostro et al. for topic 8 (**A**), topic 9 (**B**) and topic 10 (**C**).

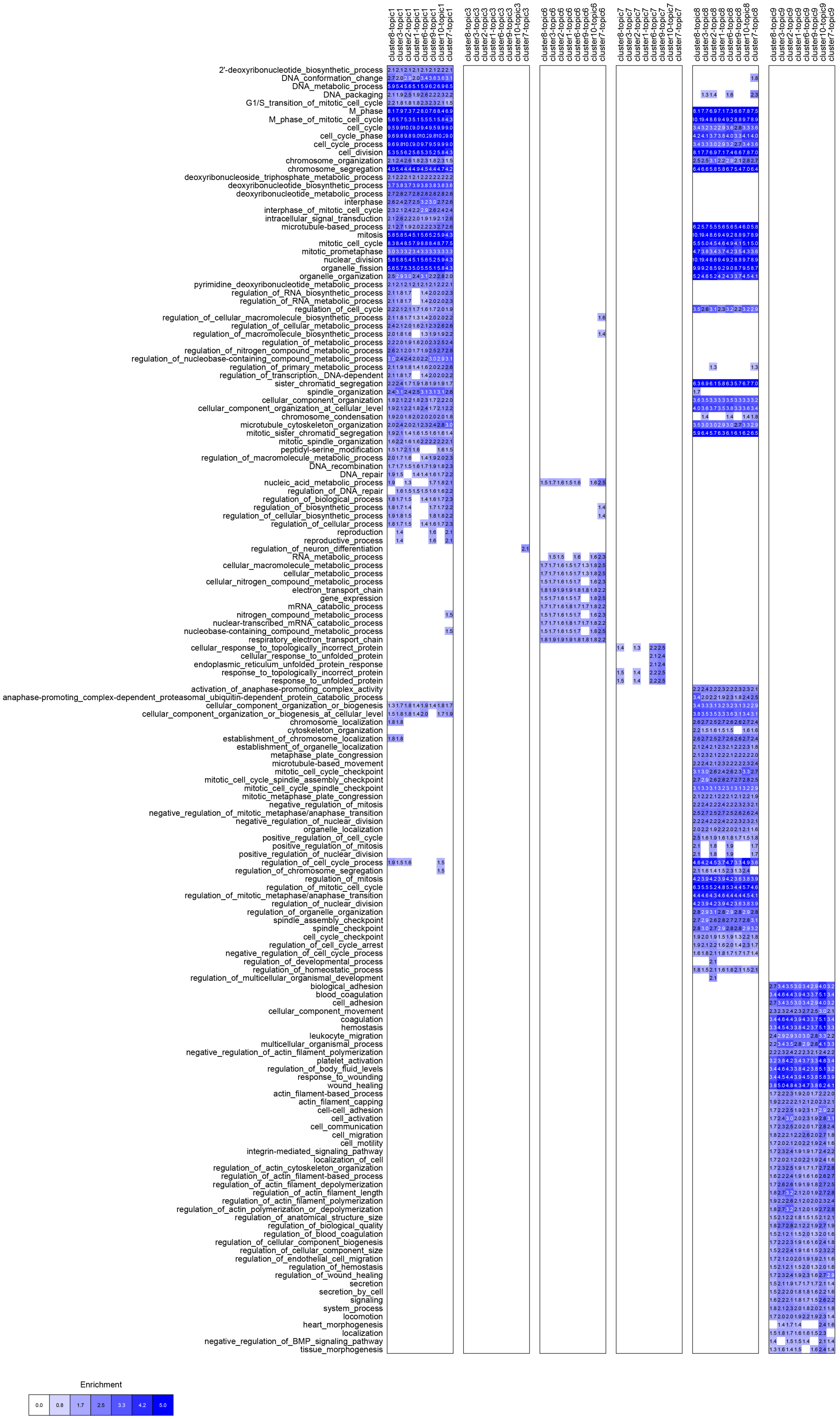

**Supplementary Figure 21.** GO enrichment for genes in each LDA topic-specific network inferred by scMTNI on the human hematopoietic differentiation data from Buenrostro et al. We used FDR $<0.01$ to determine significantly enriched terms. The blue intensity of the heat map is proportional to $-\log10(\text{FDR})$ of enrichment.

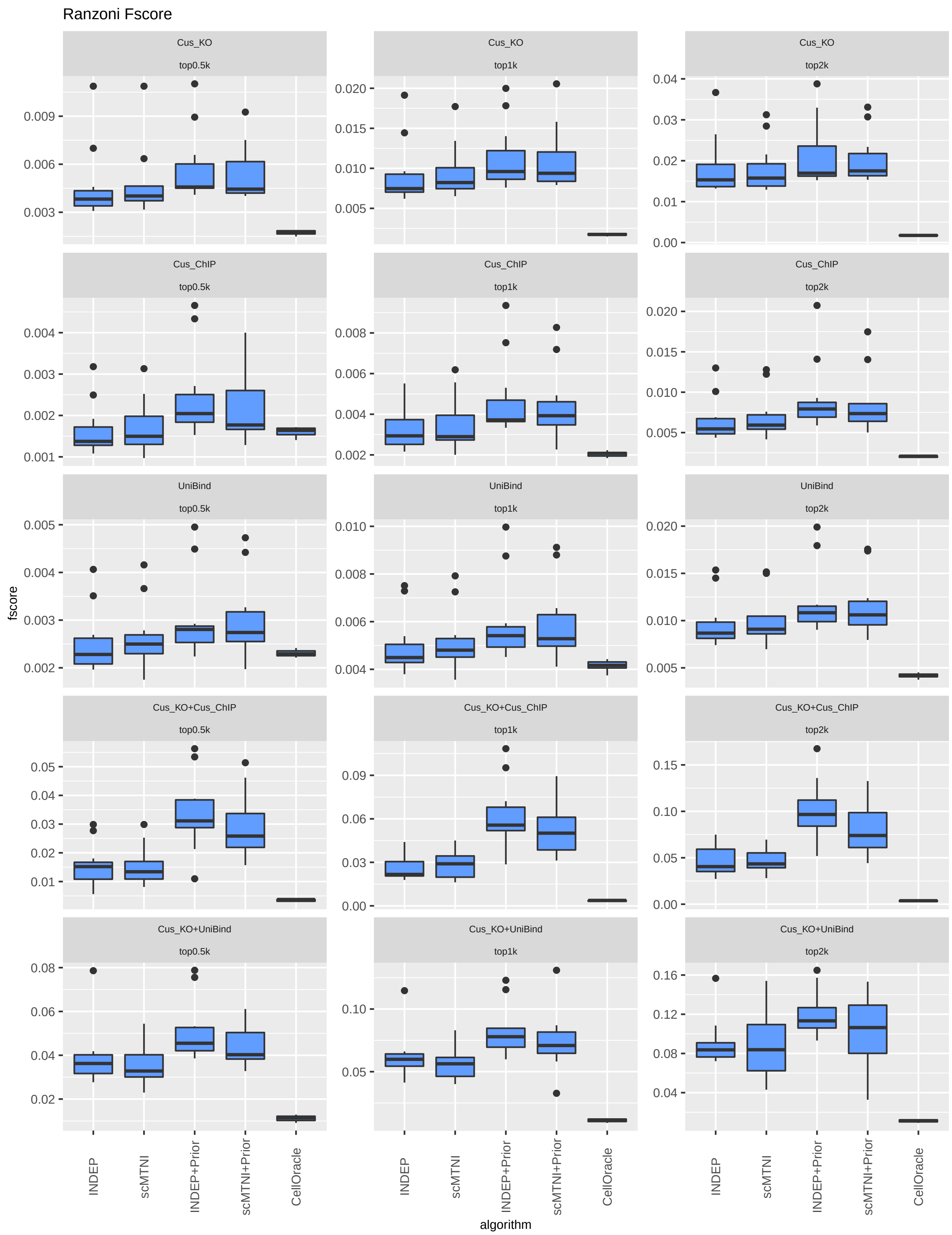

**Supplementary Figure 22.** F-score of top 500, 1k, 2k edges in predicted networks of scMTNI, scMTNI+Prior, INDEP, INDEP+Prior, and CellOracle on human fetal hematopoiesis data from Ranzoni et al. using the fine lineage. F-scores of $n = 11$ cell clusters are computed on five different gold standard datasets. In the boxplot, the horizontal middle line of each plot is the median. The bounds of the box are 0.25 quantile ($Q_1$) and 0.75 quantile ($Q_3$). The upper whisker is the minimum of the maximum value and $Q_3 + 1.5 * IQR$, where $IQR = Q_3 - Q_1$. The lower whisker is the maximum of the minimum value and $Q_1 - 1.5 * IQR$.

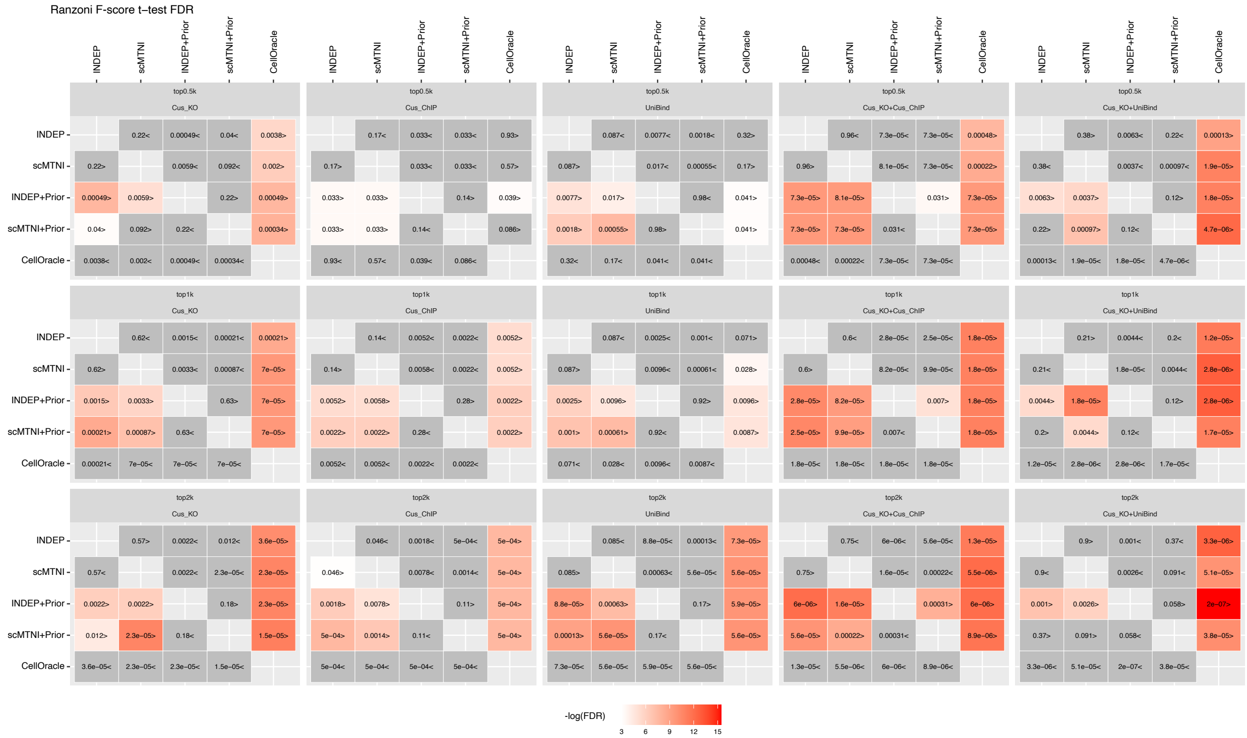

**Supplementary Figure 23.** Heatmap of FDR corrected T-test pvalues comparing the F-score of top 500, 1k, 2k edges in predicted networks of scMTNI, scMTNI+Prior, INDEP, INDEP+Prior, and CellOracle on human fetal hematopoiesis data from Ranzoni et al. using the fine lineage. Shown are relative performances on five gold standard datasets. The two-sided paired t-test is conducted on F-scores of $n = 11$ cell clusters for every pair of algorithms, comparing whether the row algorithm's F-score is higher than the column algorithm's F-score. Significant difference (FDR$<$0.05) is highlighted using a white-red colormap (the color scale for $-\log(\text{FDR})$). Non-significance is colored in gray. The sign"$<$" or "$>$" specifies whether the row algorithm's F-scores were worse or better than the column algorithm's F-scores.

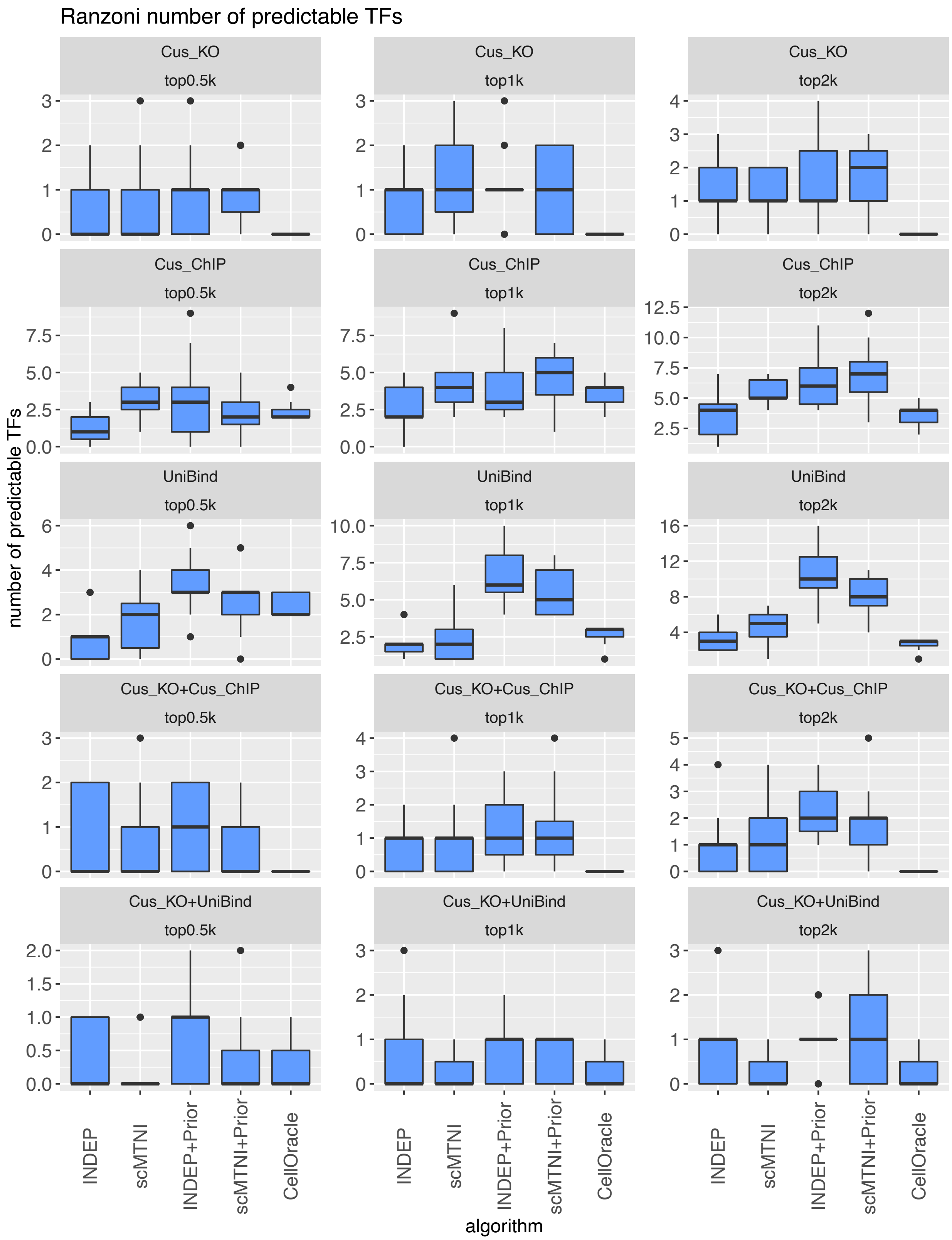

**Supplementary Figure 24.** Number of predictable TFs of $n = 11$ cell clusters in top 500, 1k, 2k edges in predicted networks of scMTNI, scMTNI+Prior, INDEP, INDEP+Prior, and CellOracle on fetal human hematopoietic differentiation data from Ranzoni et al. using the fine lineage. Each row corresponds to a particular gold standard and column corresponds to the number of top edges. In the boxplot, the horizontal middle line of each plot is the median. The bounds of the box are 0.25 quantile ($Q_1$) and 0.75 quantile ($Q_3$). The upper whisker is the minimum of the maximum value and $Q_3 + 1.5 * IQR$, where $IQR = Q_3 - Q_1$. The lower whisker is the maximum of the minimum value and $Q_1 - 1.5 * IQR$.

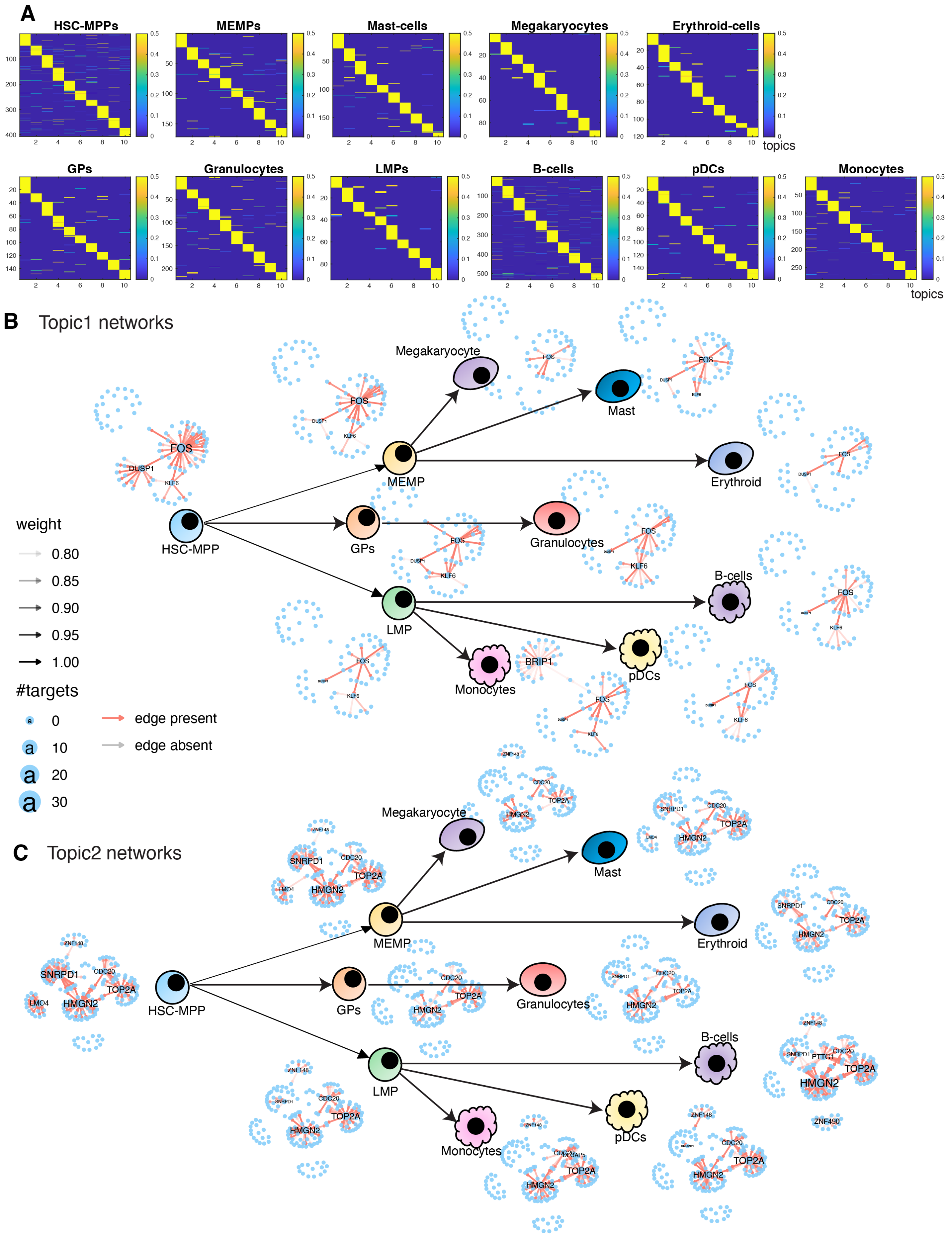

**Supplementary Figure 25.** LDA analysis for scMTNI inferred networks on human fetal hematopoiesis from Ranzoni et al. using the fine lineage structure. **A.** The document-topic weight matrix for all regulators for all cell clusters with 10 topics. Topic-specific networks across each cell cluster for topic 1 (**B**) and topic 2 (**C**).

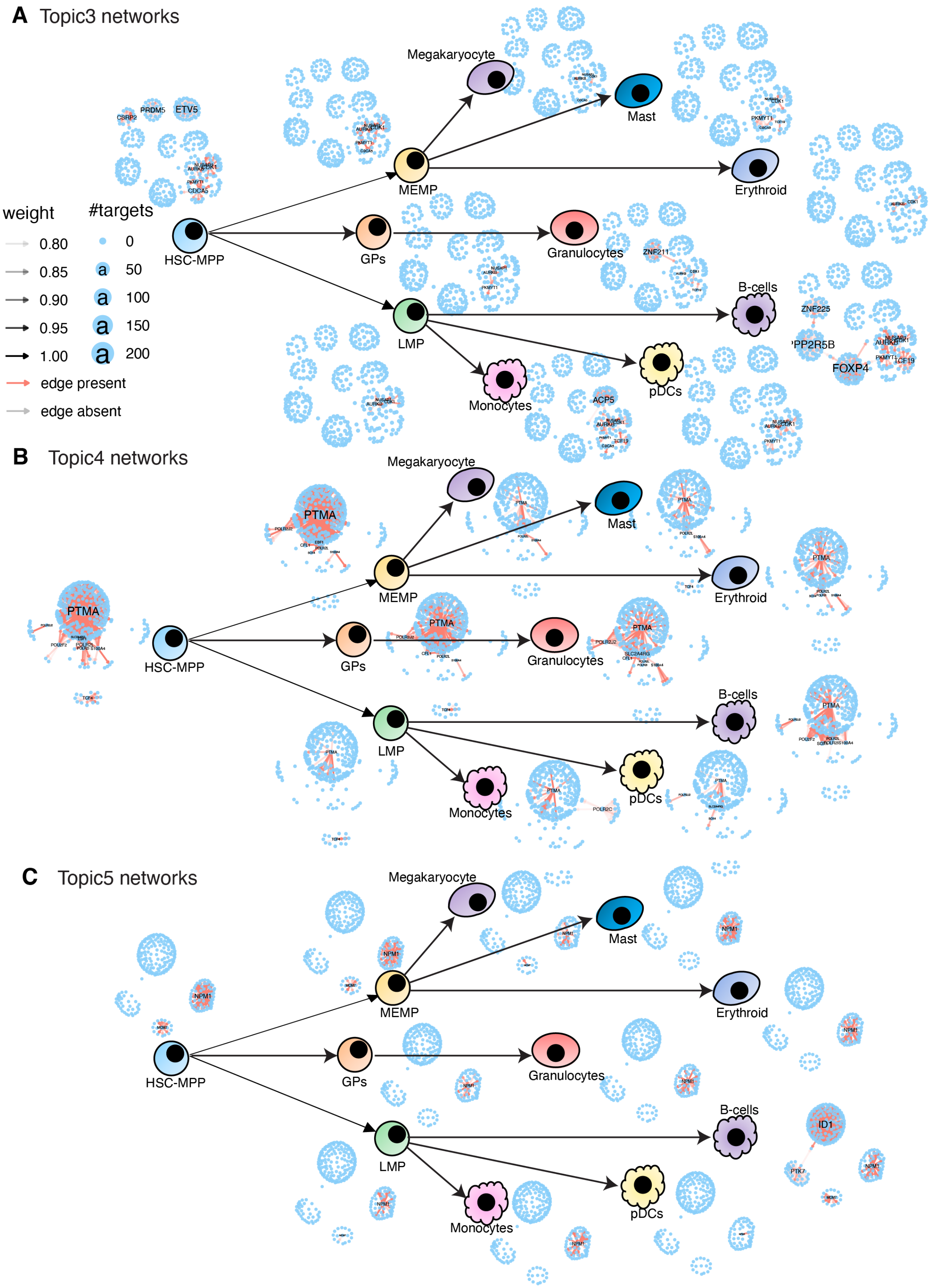

**Supplementary Figure 26.** Topic-specific networks obtained from scMTNI inferred networks on fetal human hematopoietic differentiation data from Ranzoni et al using the fine lineage. Shown are topic-specific networks for topic 3 (**A**), topic 4 (**B**) for topic 5 (**C**).

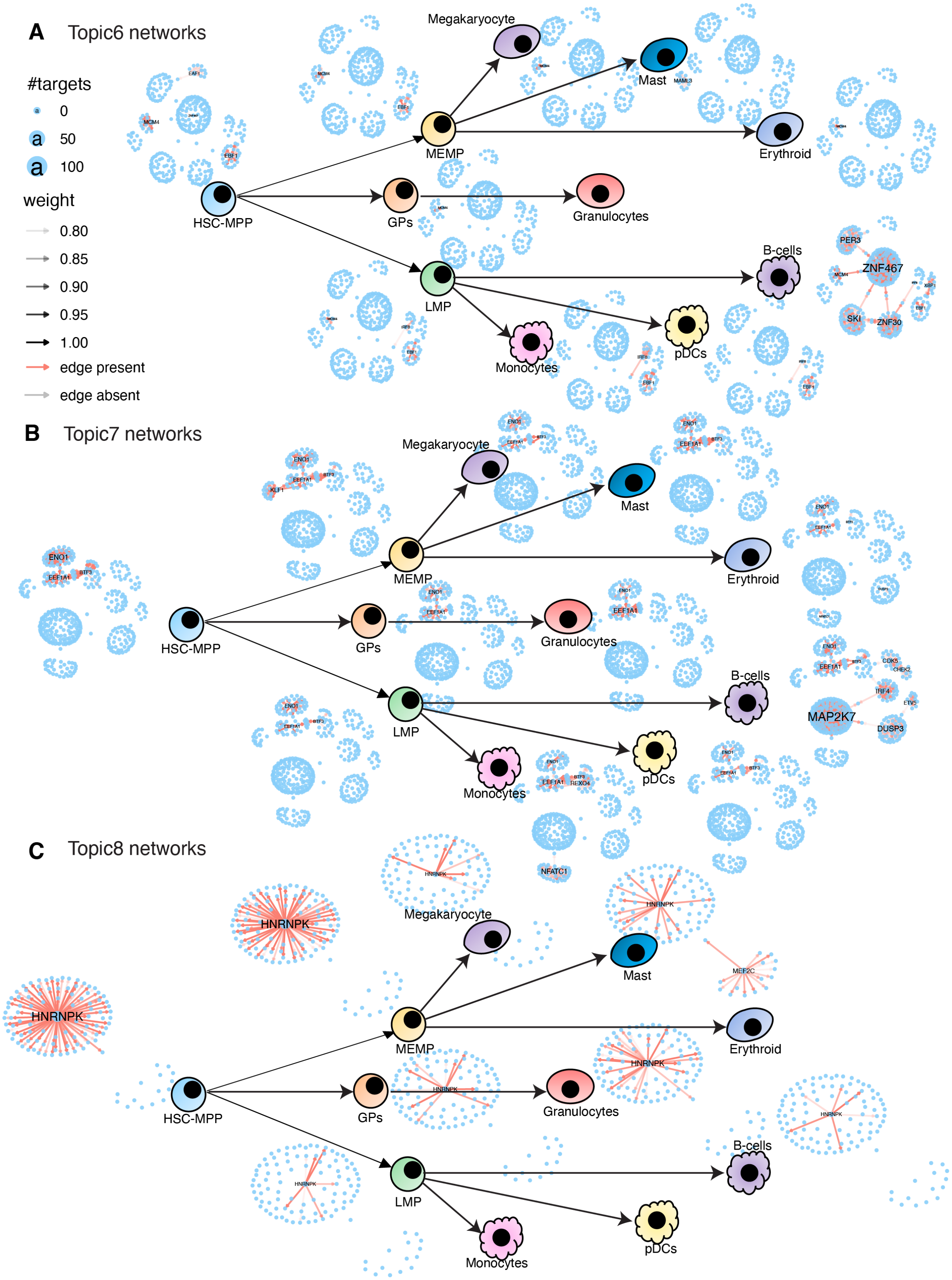

**Supplementary Figure 27.** Topic-specific networks obtained from scMTNI inferred networks on human fetal hematopoiesis data from Ranzoni et al using the fine lineage for topic 6 (**A**), topic 7 (**B**), and topic 8 (**C**).

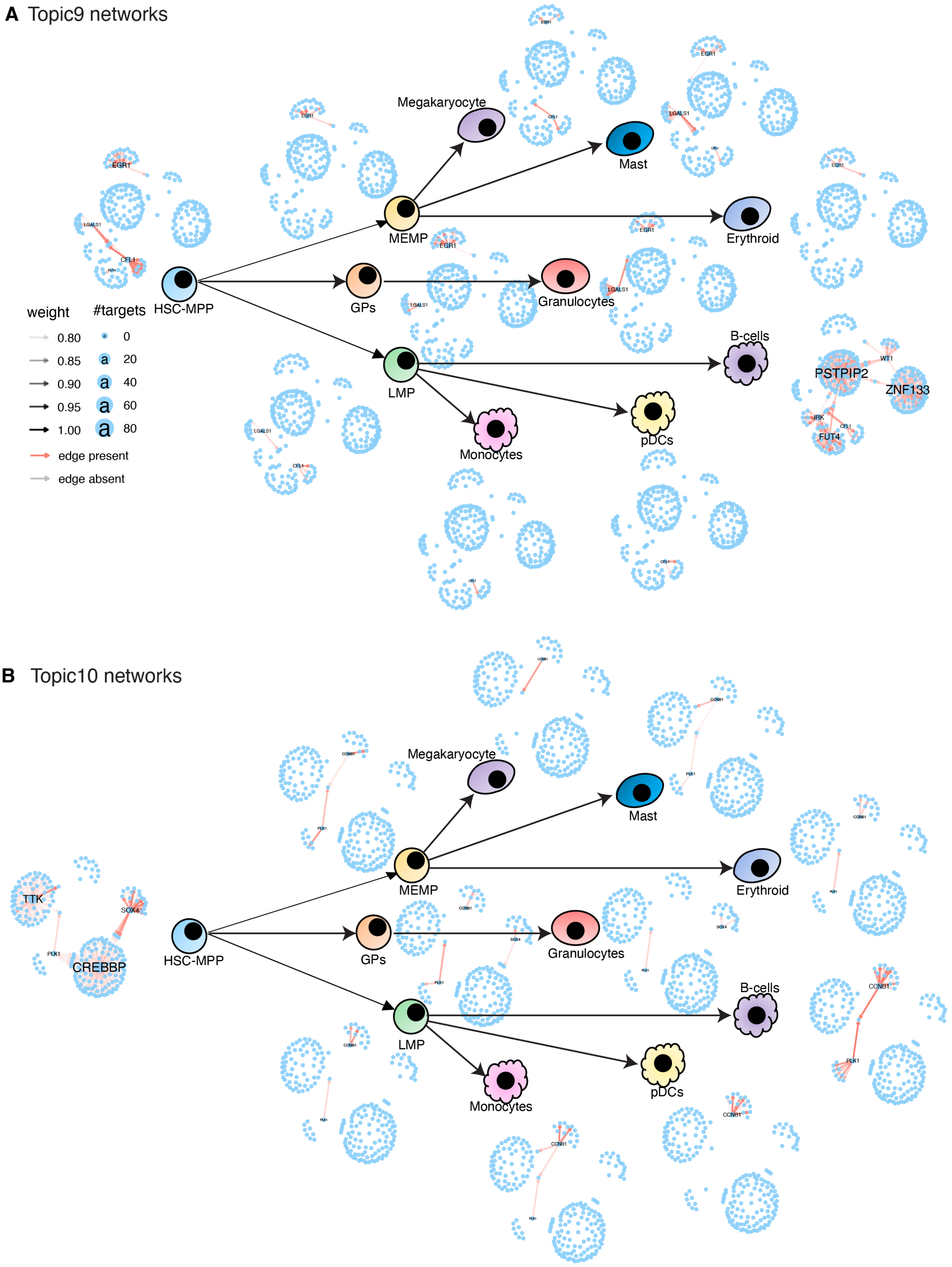

**Supplementary Figure 28.** Topic-specific networks obtained from scMTNI inferred networks on human fetal hematopoiesis data from Ranzoni et al using the fine lineage for topic 9 (**A**) and topic 10 (**B**).

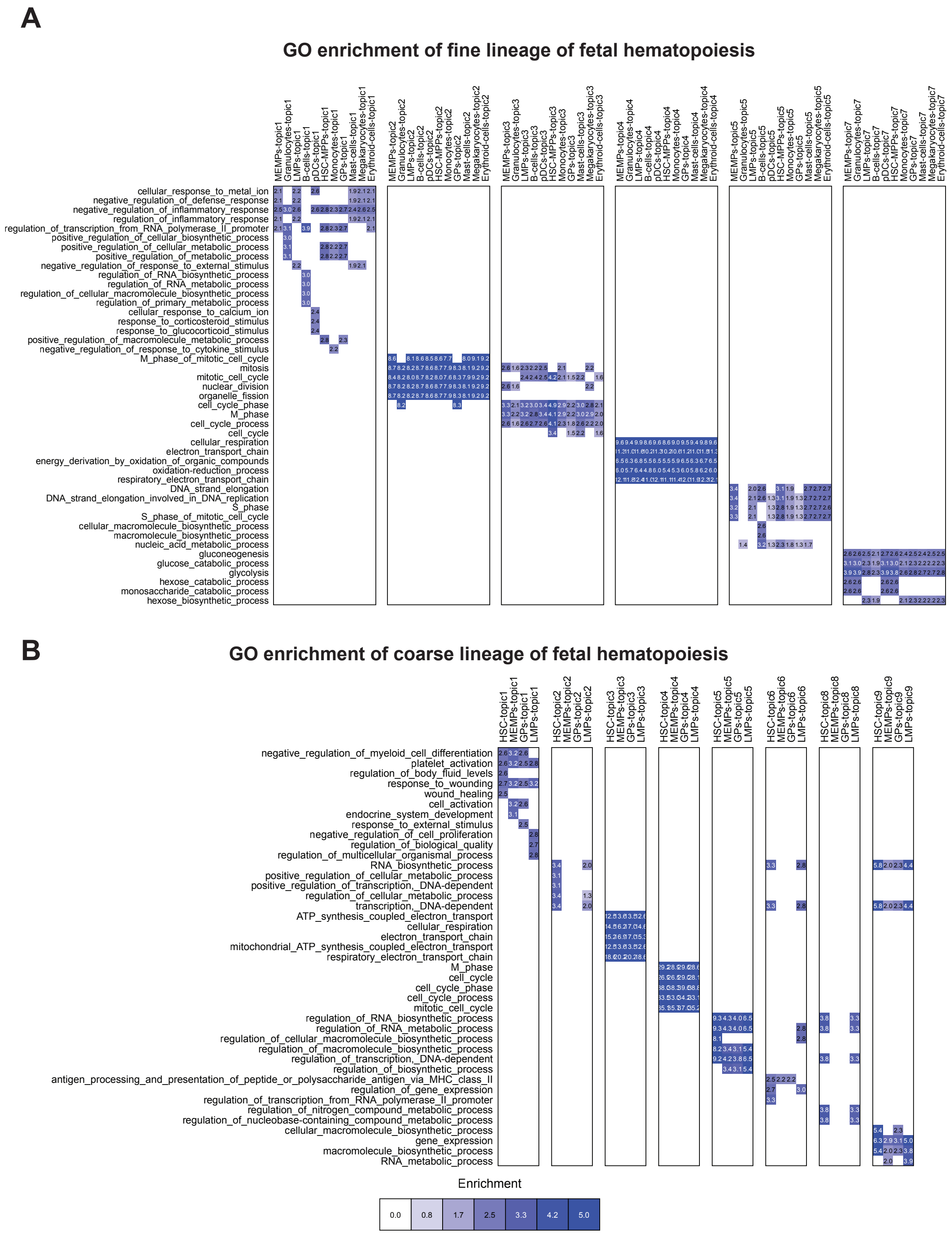

**Supplementary Figure 29.** Gene Ontology enrichment of LDA topics in fetal hematopoiesis from Ranzoni et al. **A.** Gene ontology terms enriched in genes for each topic identified by LDA for the fine lineage. Top 5 terms for each topic and cell type, with an FDR<0.01 are shown. **B.** Gene ontology terms enriched in genes for each topic identified by LDA for the coarse lineage. Top 5 terms for each topic and cell type, with an FDR<0.01 are shown. For both **A** and **B**, the blue intensity of the heat map is proportional to $-\log10(\text{FDR})$ of enrichment.

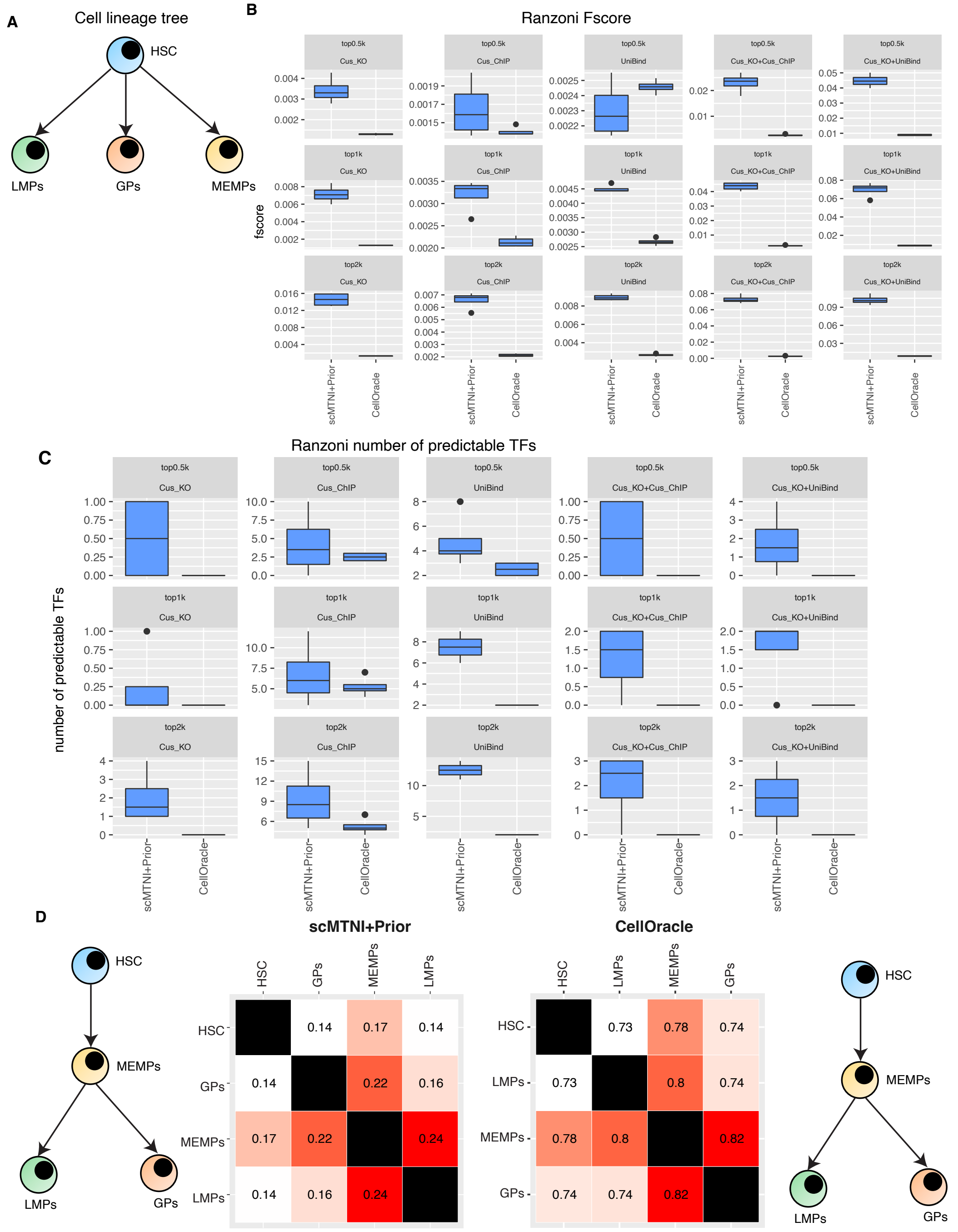

**Supplementary Figure 30.** Application of scMTNI+Prior and CellOracle to human fetal hematopoiesis data from Ranzoni et al using coarse lineage structure. **A.** The cell lineage structure used as input to scMTNI. **B.** Performance of scMTNI and CellOracle assessed using F-score of $n = 4$ cell clusters on five different gold standards (rows). Each column shows values for the top 500, 1k or 2k edges. **C.** Number of predictable TFs of $n = 4$ cell clusters for scMTNI+Prior and CellOracle for five different gold standards using the top 500, 1k or 2k edges. In both **B** and **C**, the horizontal middle line of each plot is the median. The bounds of the box are 0.25 quantile ($Q_1$) and 0.75 quantile ($Q_3$). The upper whisker is the minimum of the maximum value and $Q_3 + 1.5 * IQR$, where $IQR = Q_3 - Q_1$. The lower whisker is the maximum of the minimum value and $Q_1 - 1.5 * IQR$. **D.** Inferred cell lineage trees using minimum spanning trees on the pairwise network similarity. The inferred cell-lineage tree for scMTNI+Prior is shown on the left, and inferred cell-lineage tree for CellOracle is shown on the right. The heatmap shows the pairwise similarity of networks from each cell cluster assessed using F-score on the top 5k edges.

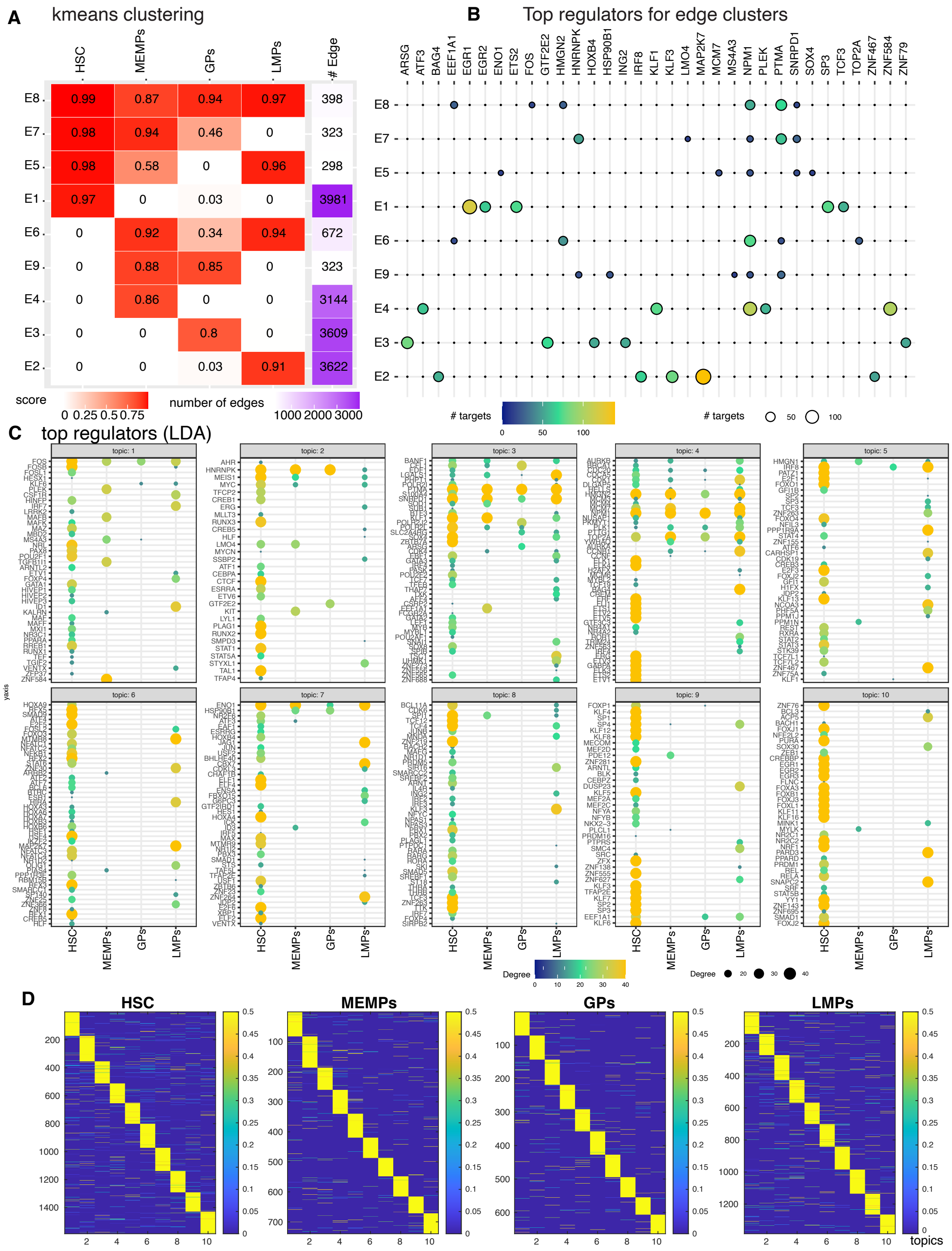

**Supplementary Figure 31.** K-means and LDA analysis of scMTNI+Prior inferred networks on the human fetal hematopoiesis data from Ranzoni et al using the coarse lineage structure. **A.** k-means analysis of scMTNI inferred networks showing edge cluster patterns. **B.** The top regulators associated with each edge cluster. **C.** LDA analysis on the scMTNI inferred networks. Top regulators for each topic obtained from LDA analysis of scMTNI inferred networks. **D.** The document-topic weight matrix for all regulators for all cell clusters with 10 topics.

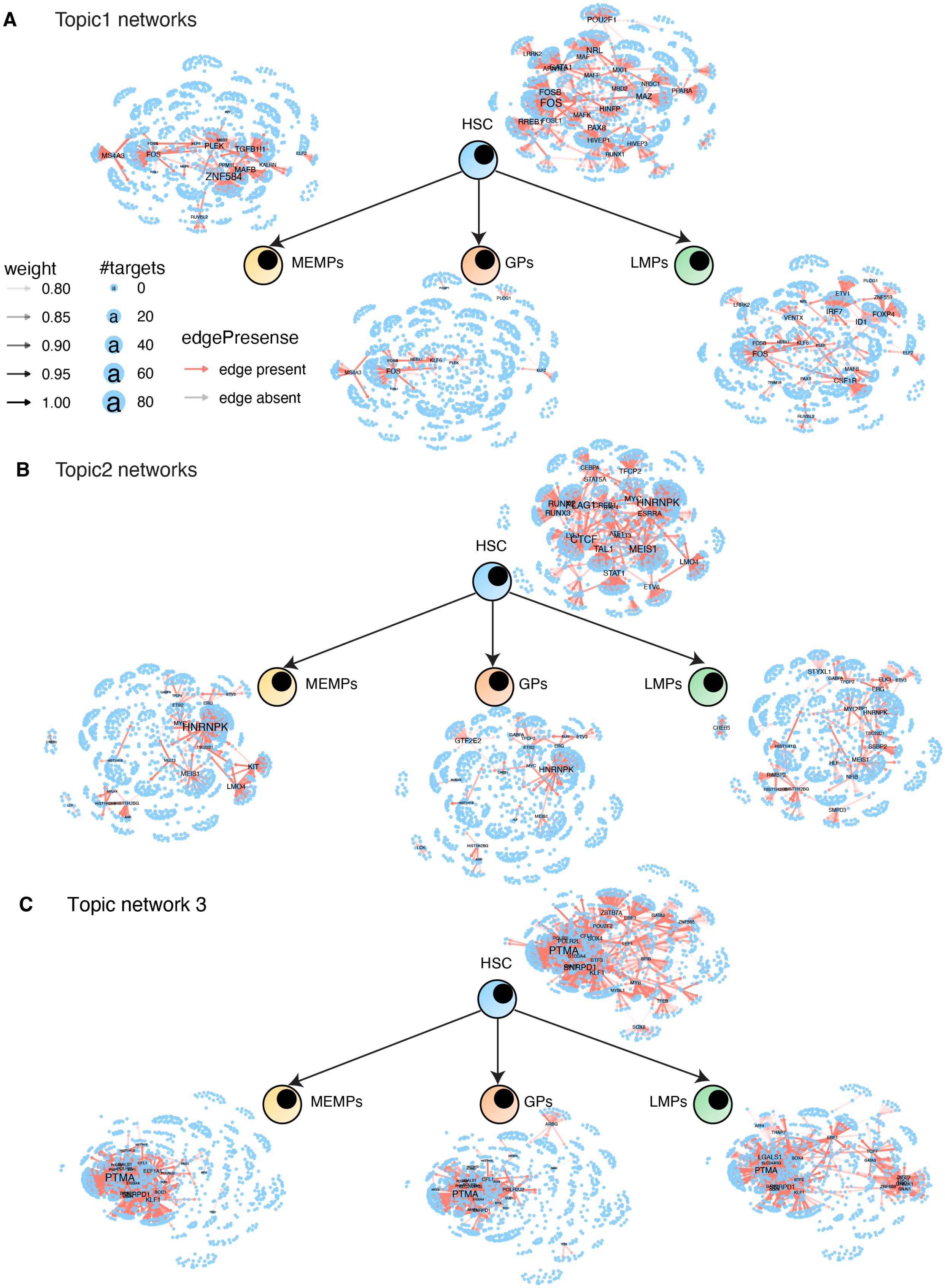

**Supplementary Figure 32.** Topic-specific networks obtained from scMTNI-inferred networks for human fetal hematopoiesis data with coarse tree from Ranzoni et al for topic 1 (**A**), topic 2 (**B**) and topic 3 (**C**).

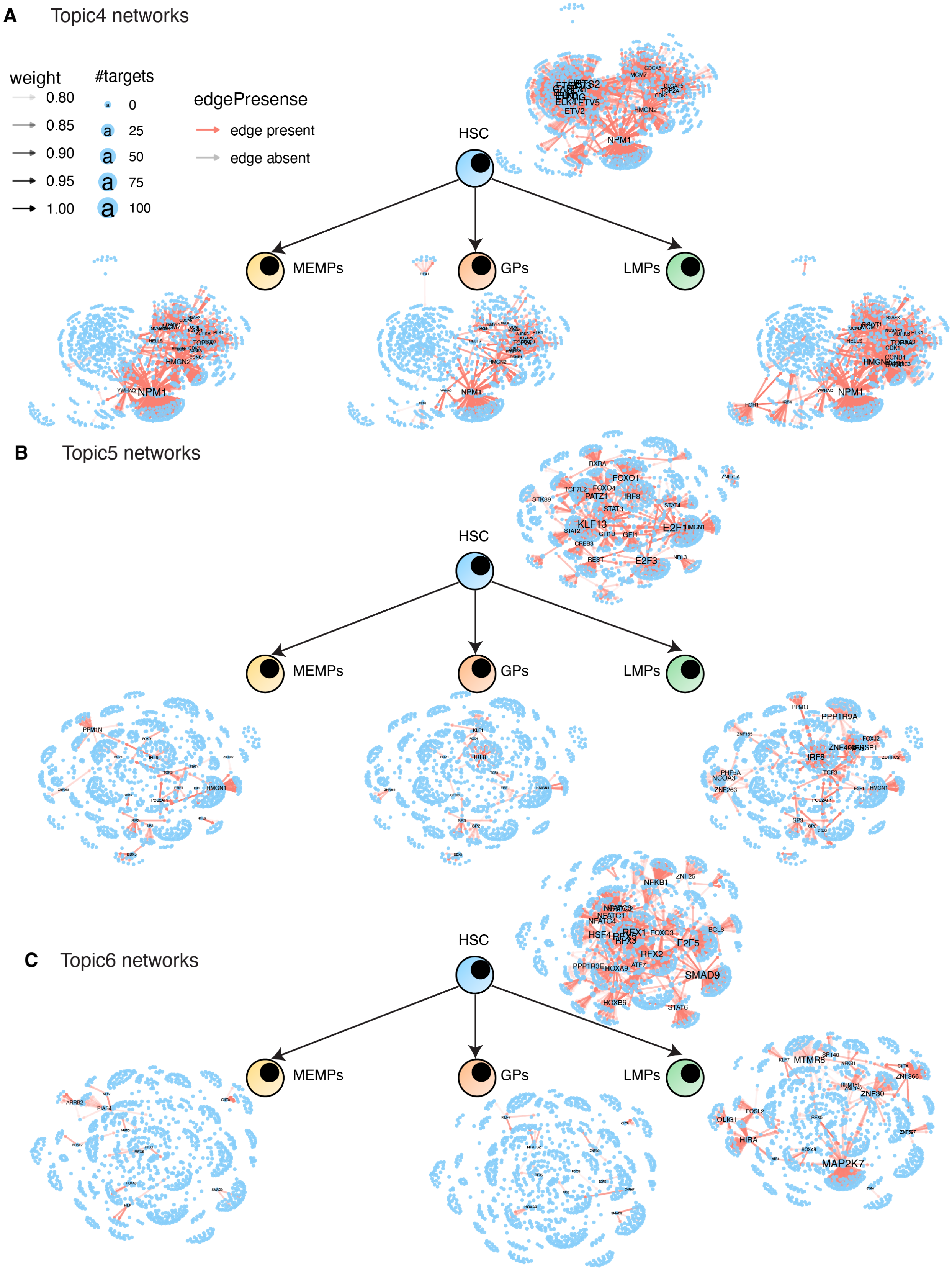

**Supplementary Figure 33.** Topic-specific networks obtained from scMTNI-inferred networks for human fetal hematopoiesis data with coarse tree from Ranzoni et al for topic 4 (**A**), topic 5 (**B**) and topic 6 (**C**).

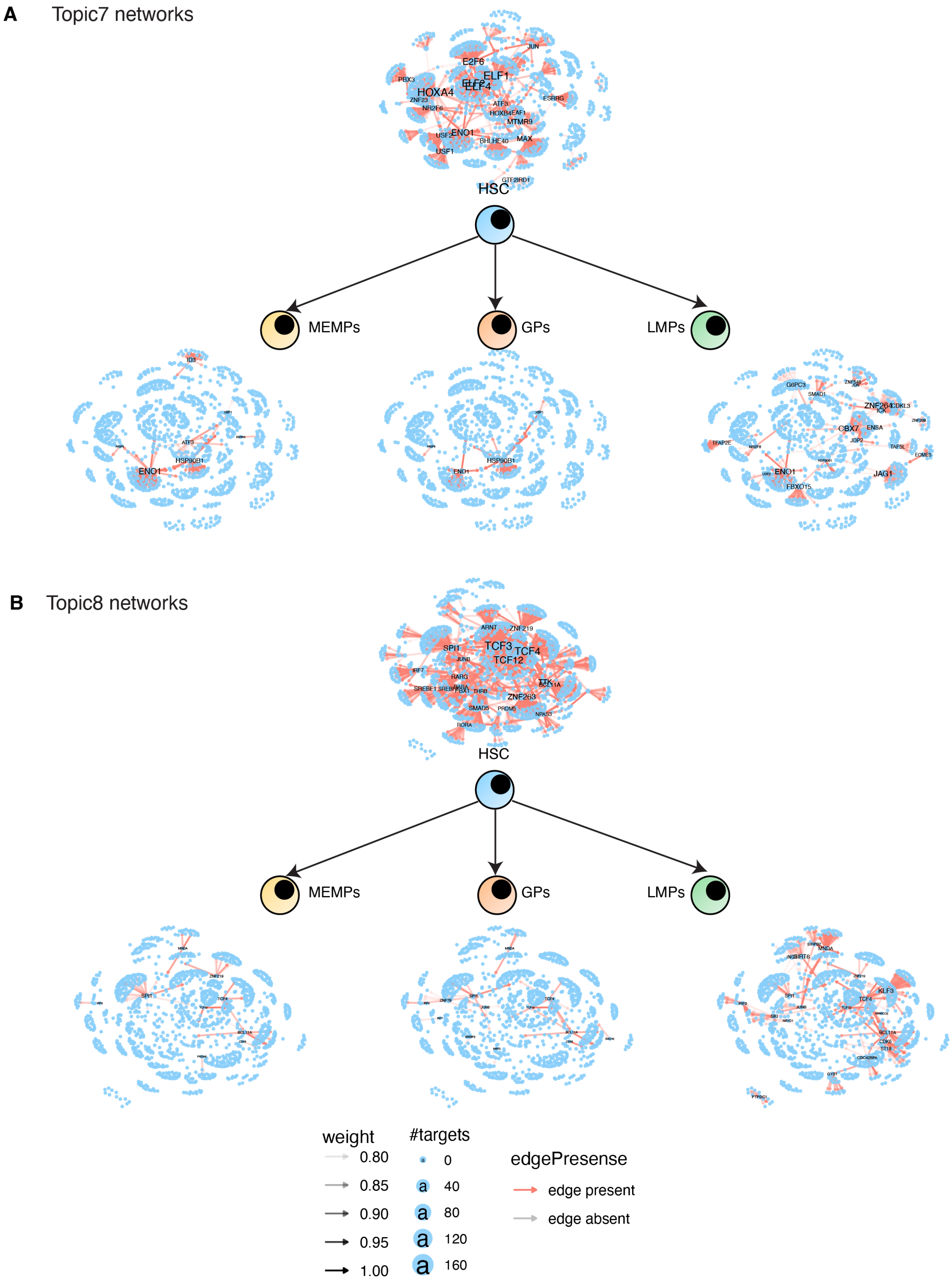

**Supplementary Figure 34.** Topic-specific networks obtained from scMTNI-inferred networks for human fetal hematopoiesis data with coarse tree from Ranzoni et al for topic 7 (**A**) and topic 8 (**B**).

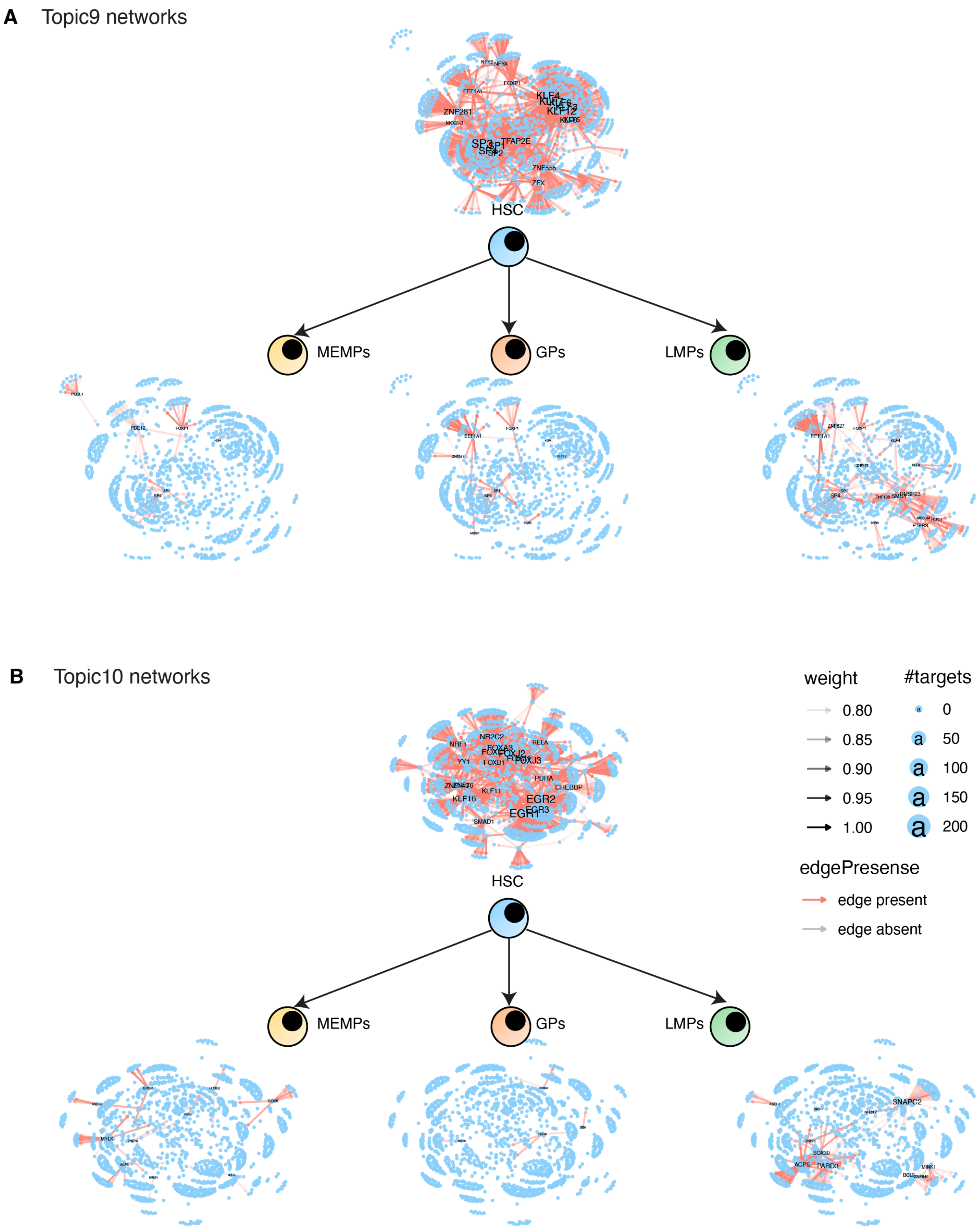

**Supplementary Figure 35.** Topic-specific networks obtained from scMTNI-inferred networks for human fetal hematopoiesis data with coarse tree from Ranzoni et al for topic 9 (**A**) and topic 10 (**B**).

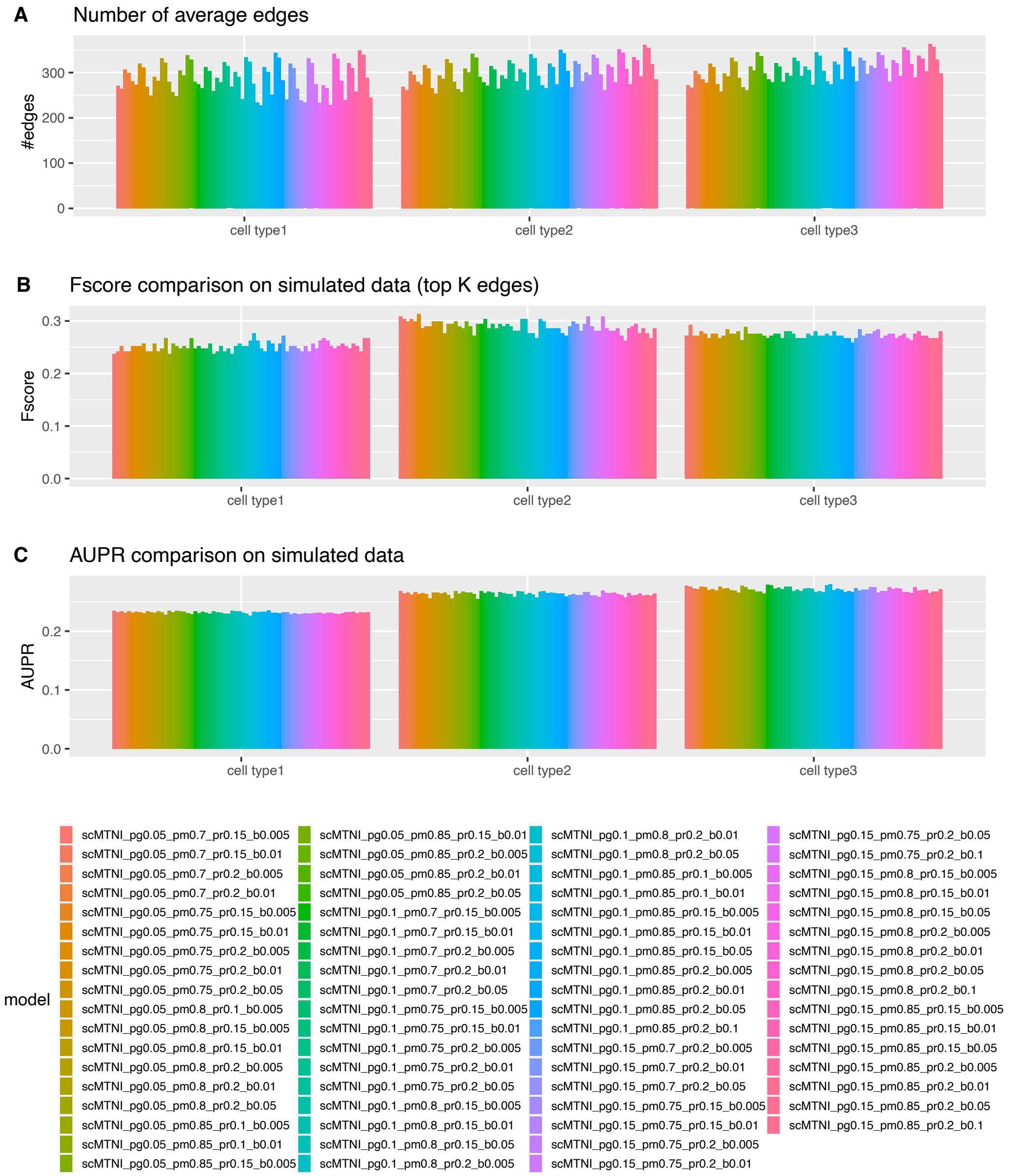

**Supplementary Figure 36.** Performance of scMTNI on simulated dataset 1 with different parameter settings of $p_r$, $p_g$, $p_m$, $b = |\beta_0|$. **A.** The average of number of edges in the inferred network across 50 subsamples. **B.** F-score comparing top edges in inferred networks to simulated ground truth networks. **C.** AUPR comparing inferred networks to simulated ground truth networks.

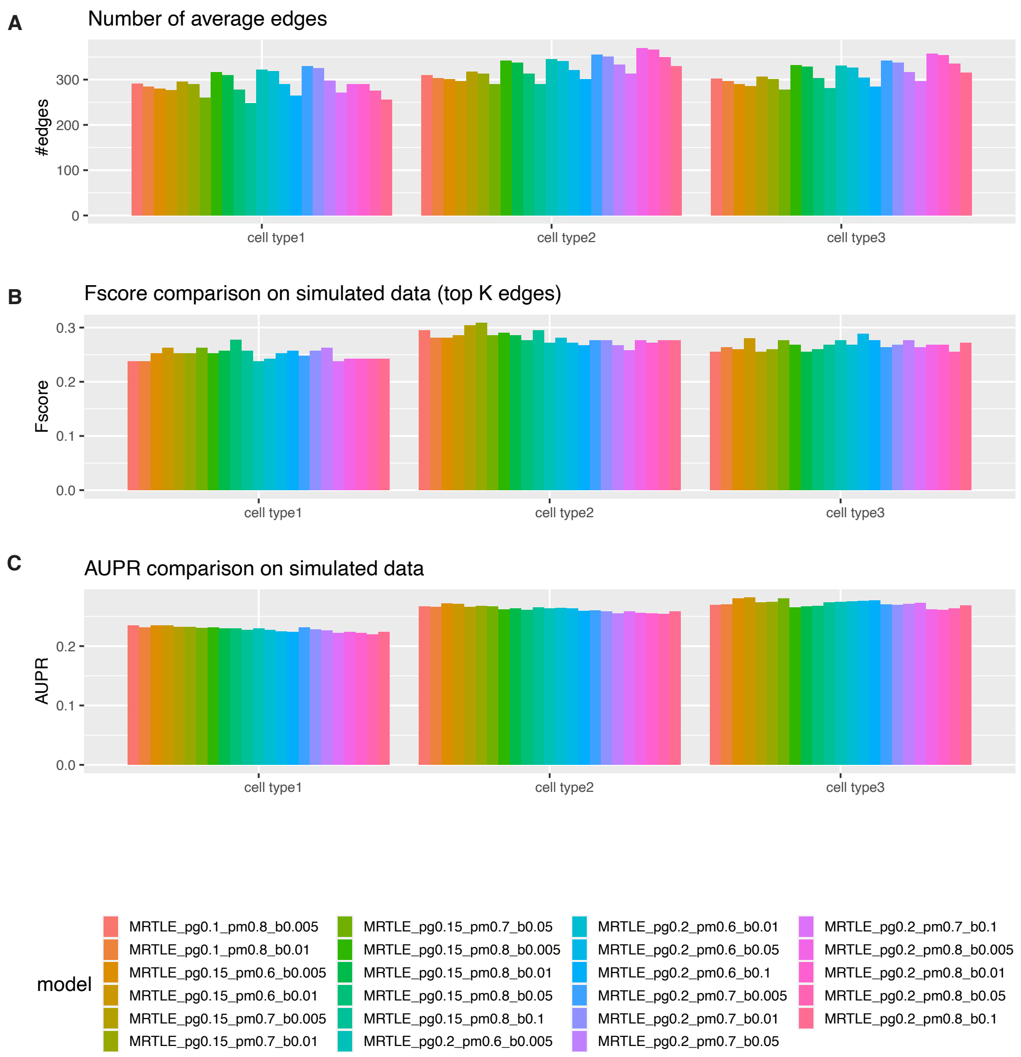

**Supplementary Figure 37.** Performance of MRTLE on simulated dataset 1 with different parameter settings of $p_g$, $p_m$, $b = |\beta_0|$. **A.** The average of number of edges output from each algorithm run across 50 subsamples. **B.** F-score comparing top edges in inferred networks to simulated ground truth networks. **C.** AUPR comparing inferred networks to ground truth networks.

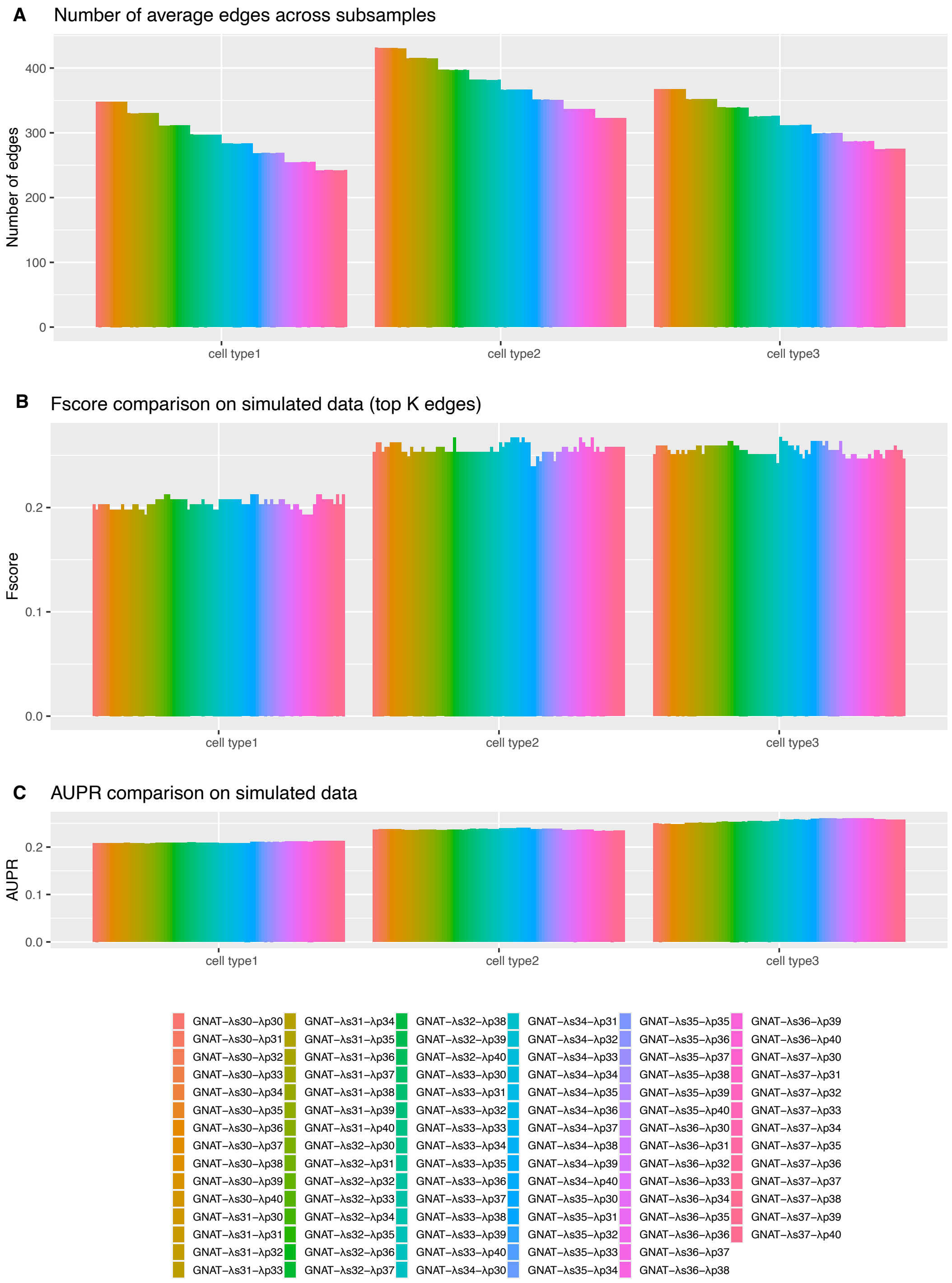

**Supplementary Figure 38.** Performance of GNAT on simulated dataset 1 with different parameter settings of $\lambda_s$ and $\lambda_p$. **A.** The average of number of edges output from each algorithm run across 50 subsamples. **B.** F-score comparing top edges in inferred networks to simulated ground truth networks. **C.** AUPR comparing inferred networks to simulated ground truth networks.

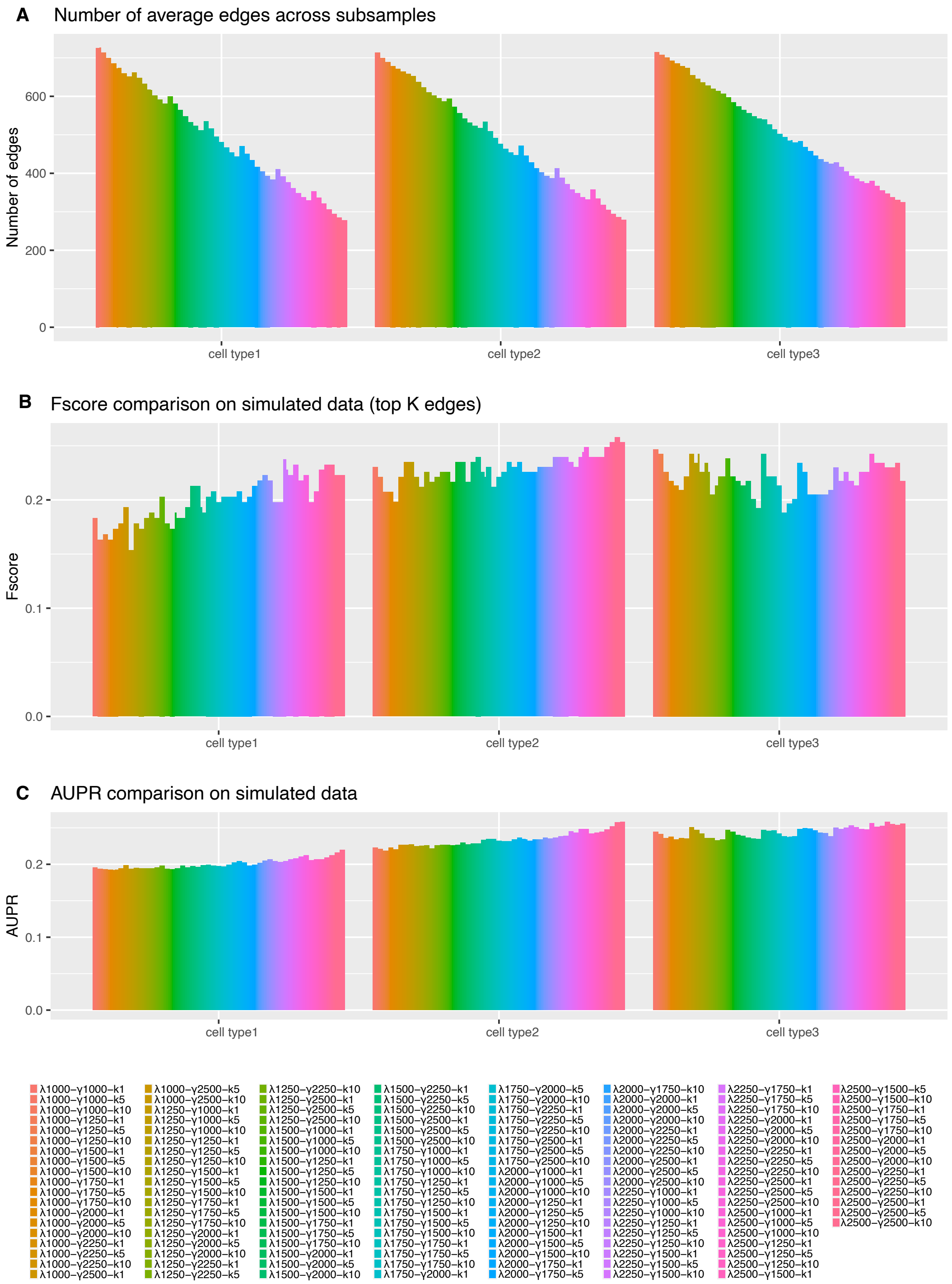

**Supplementary Figure 39.** Performance of Ontogenet on simulated dataset 1 with different parameter settings of $\lambda$, $\gamma$ and $\kappa$. **A.** The average of number of edges output from each algorithm run across 50 subsamples. **B.** F-score comparing top edges in inferred networks to simulated ground truth networks. **C.** AUPR comparing inferred networks to simulated ground truth networks.

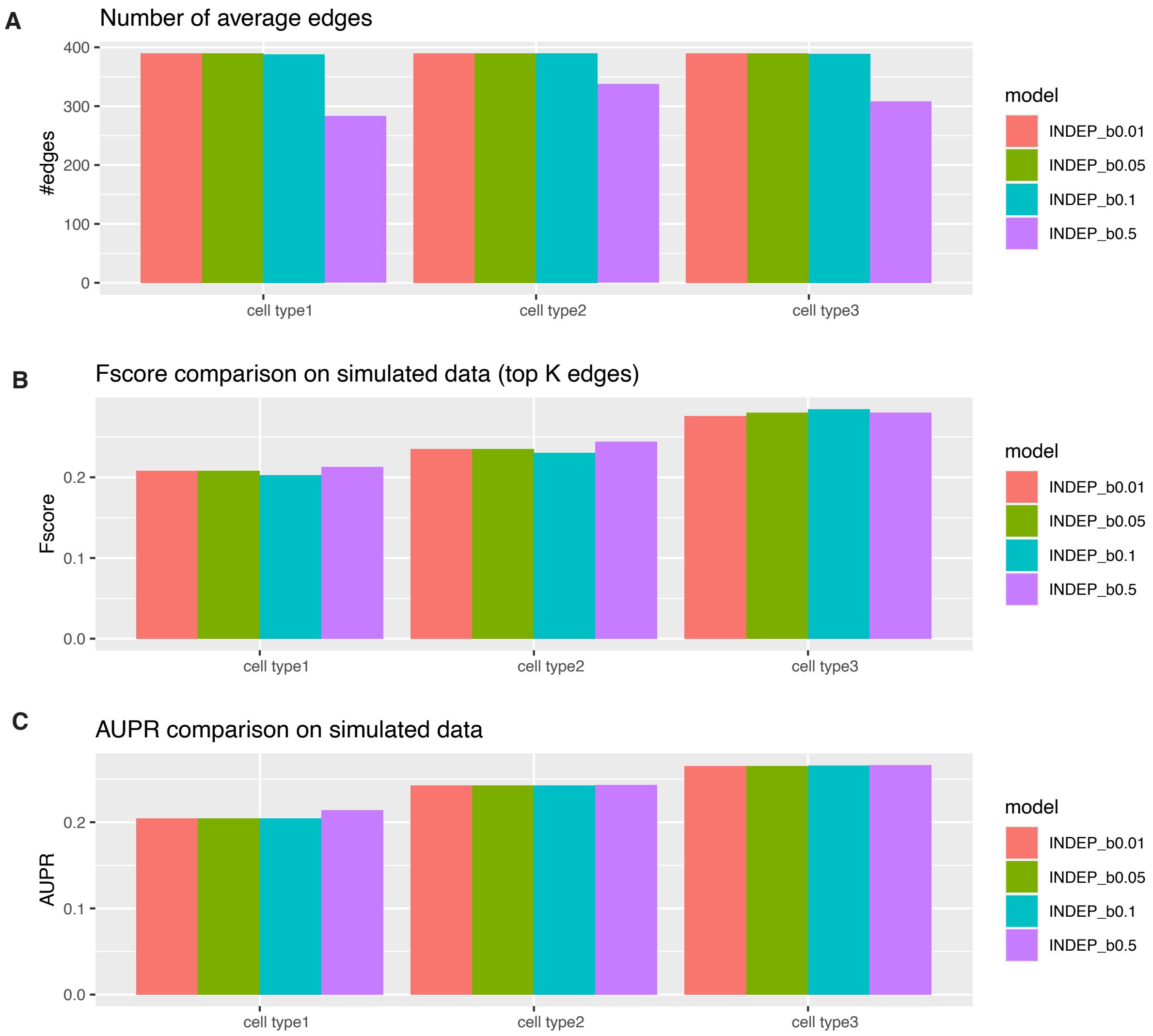

**Supplementary Figure 40.** Performance of INDEP on simulated dataset 1 with different parameter settings of $b = |\beta_0|$. **A.** The average of number of edges output from each algorithm run across 50 subsamples. **B.** F-score comparing top edges in inferred networks to simulated ground truth networks. **C.** AUPR comparing inferred networks to simulated ground truth networks.

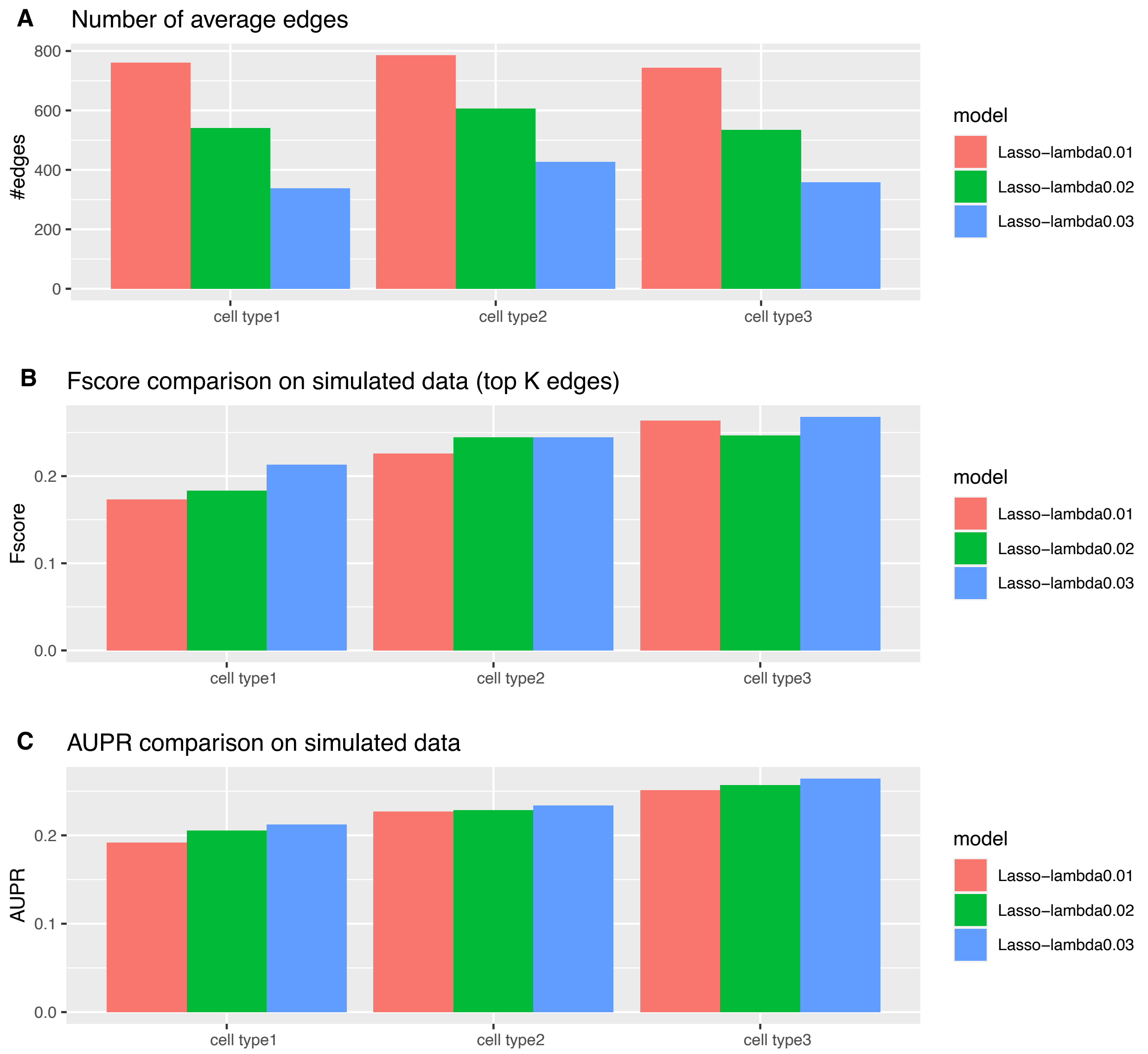

**Supplementary Figure 41.** Performance of LASSO on simulated dataset 1 with different parameter settings of $\lambda$. **A.** The average of number of edges output from each algorithm run across 50 subsamples. **B.** F-score comparing top edges in inferred networks to simulated ground truth networks. **C.** AUPR comparing inferred networks to simulated ground truth networks.

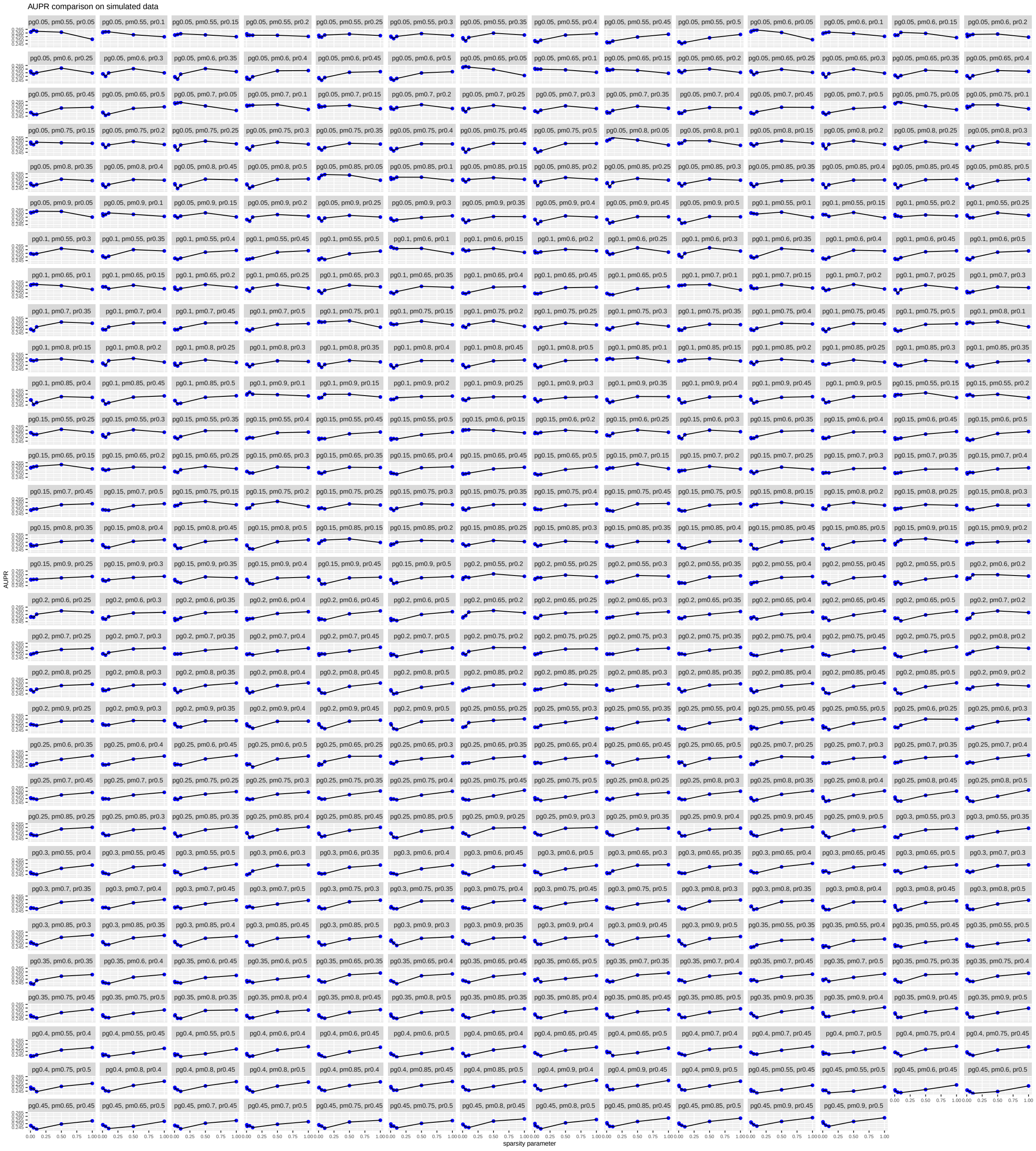

**Supplementary Figure 42.** Effect of sparsity parameter on the overall AUPR. Each subplot shows the AUPR as a function of changing the sparsity $\beta_0$ parameter (x-axis, $\beta_0 < 0$, $|\beta_0|$ is shown), while keeping the other parameters, $p_r$, $p_m$, $p_g$ fixed. The specific setting for these parameters are in the title. Shown is the average performance across 3 cell types based on AUPR on simulated dataset 1.

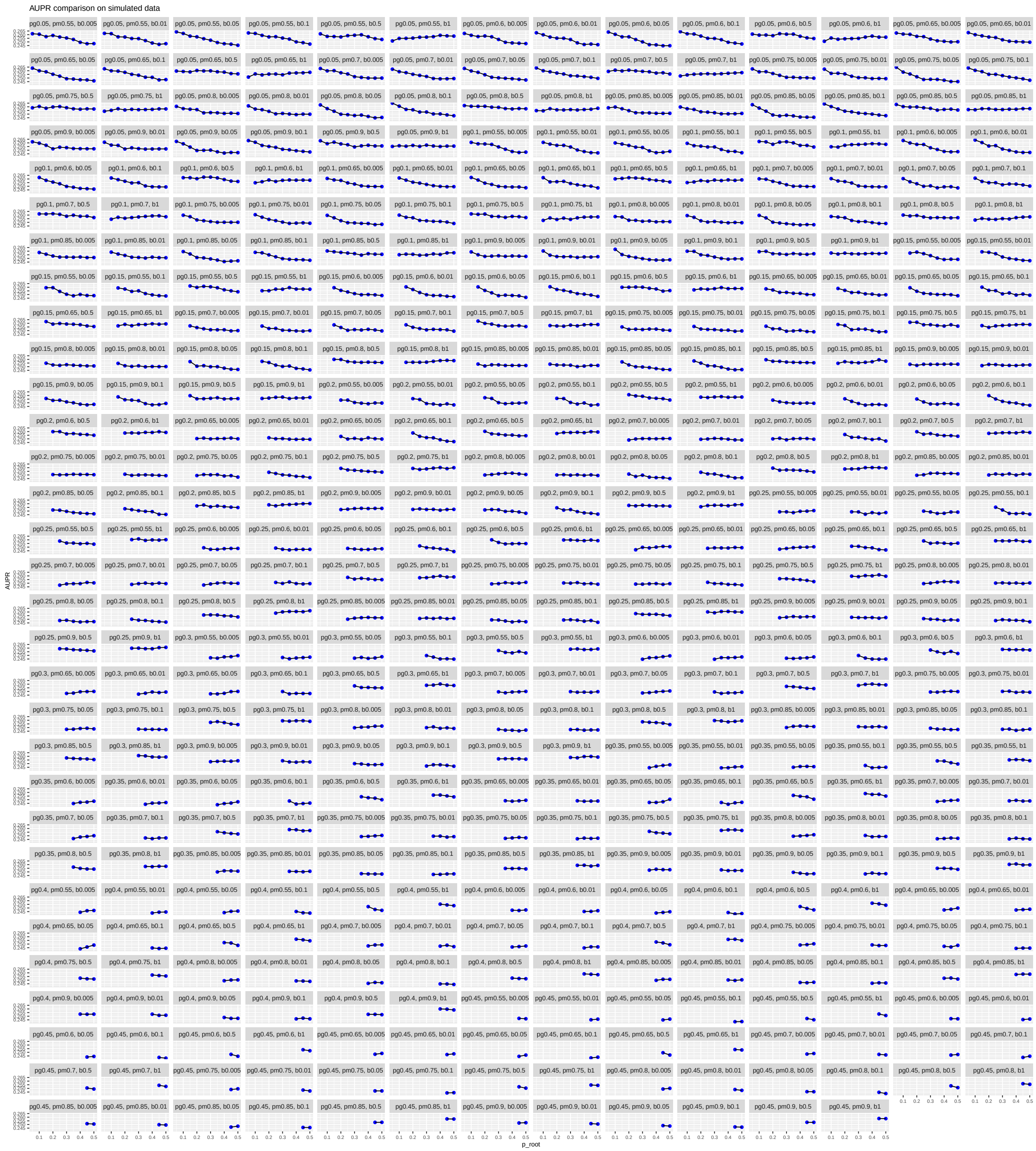

**Supplementary Figure 43.** Effect of $p_r$ parameter on the overall AUPR. Changing the $p_r$ parameter while keeping the other parameters fixed, the average performance across 3 cell types based on AUPR on simulated dataset 1.

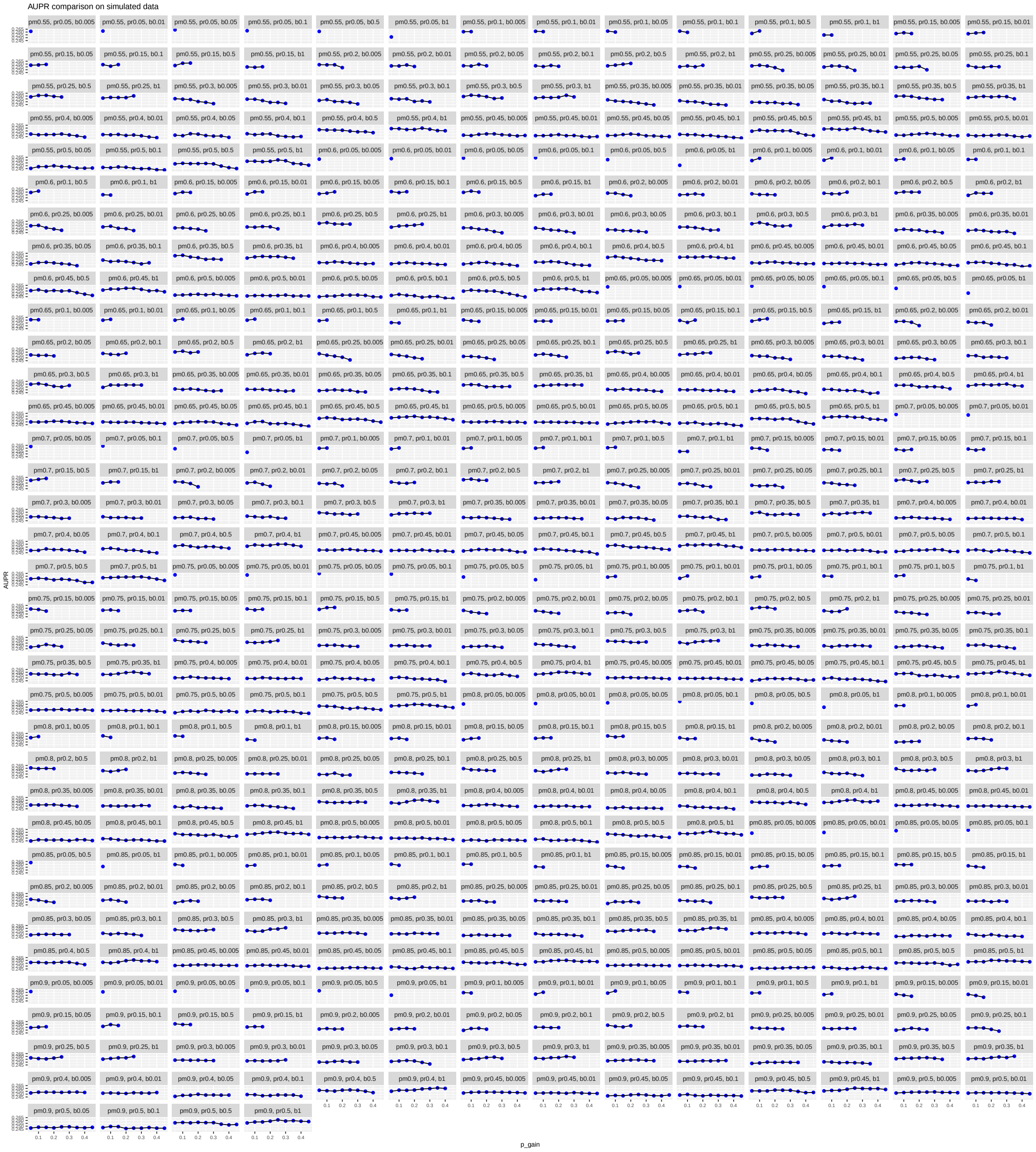

**Supplementary Figure 44.** Effect of $p_g$ parameter on the overall AUPR. Changing the $p_g$ parameter while keeping the other parameters fixed, the average performance across 3 cell types based on AUPR on simulated dataset 1.

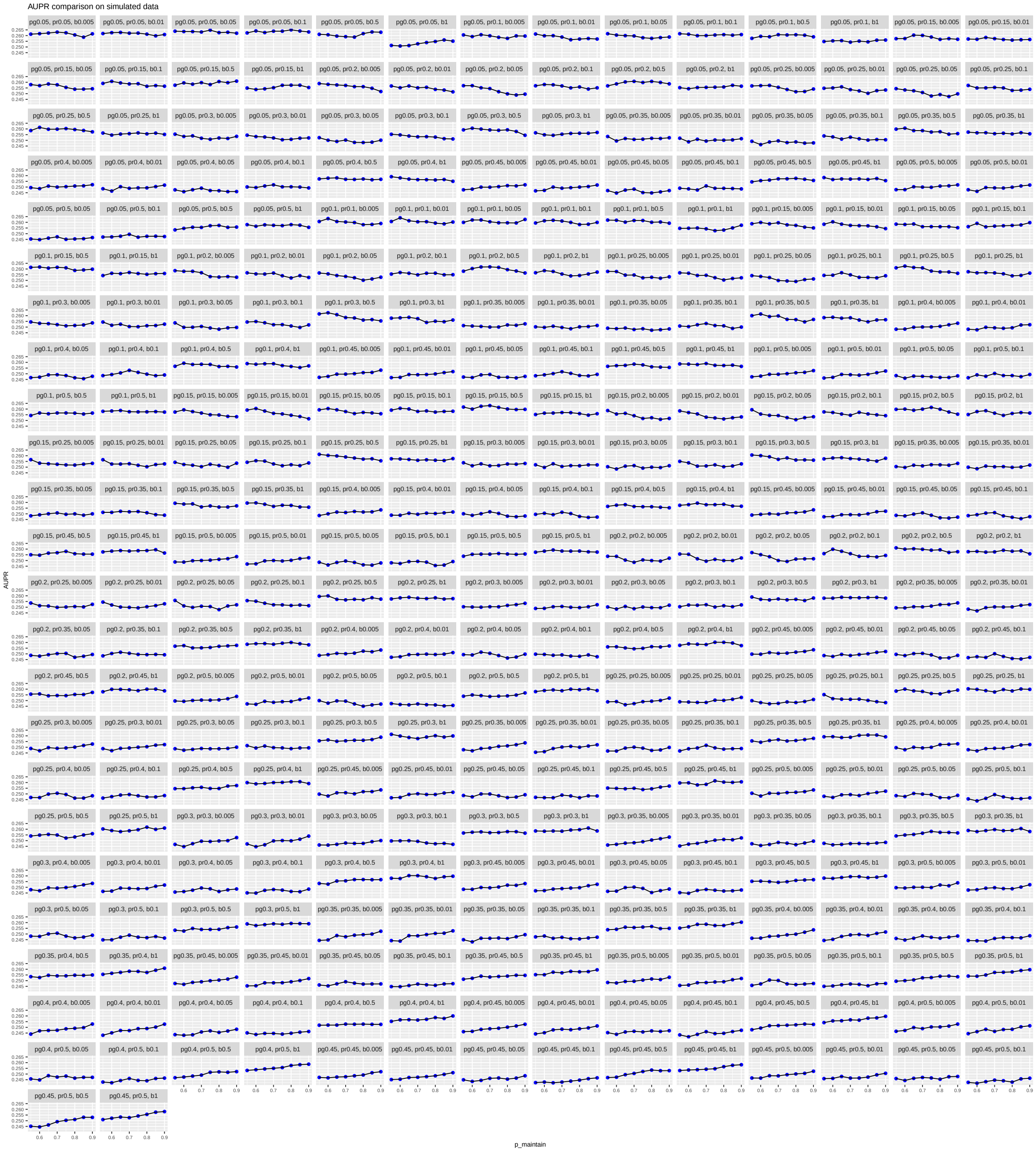

**Supplementary Figure 45.** Effect of $p_m$ parameter on the overall AUPR. Changing the $p_m$ parameter while keeping the other parameters fixed, the average performance across 3 cell types based on AUPR on simulated dataset 1.

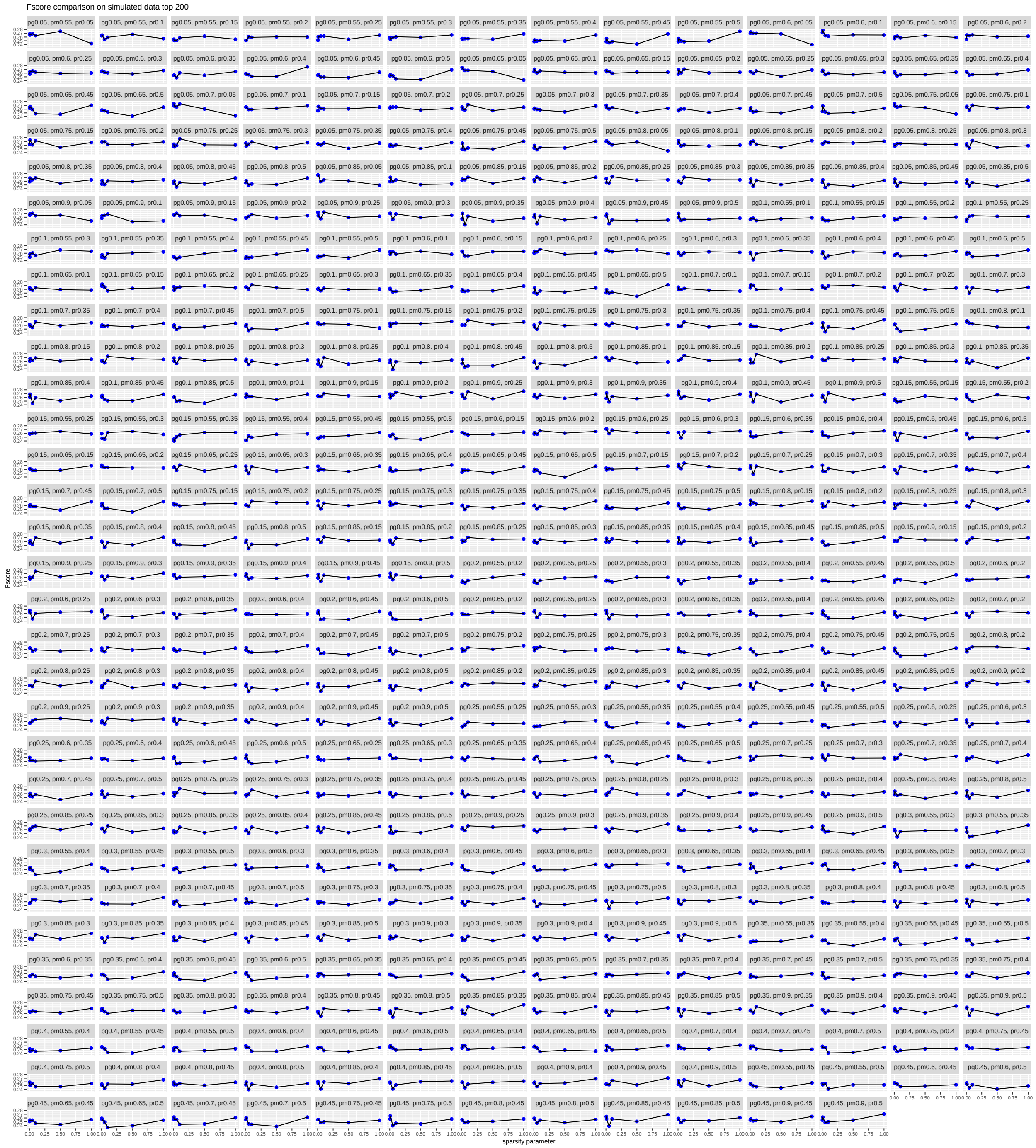

**Supplementary Figure 46.** Effect of sparsity parameter on the overall F-score. Changing the sparsity $\beta_0$ (x-axis, $\beta_0 < 0$, $|\beta_0|$ is shown) parameter while keeping the other parameters fixed, the average performance across 3 cell types based on F-score of top 200 edges on simulated dataset 1.

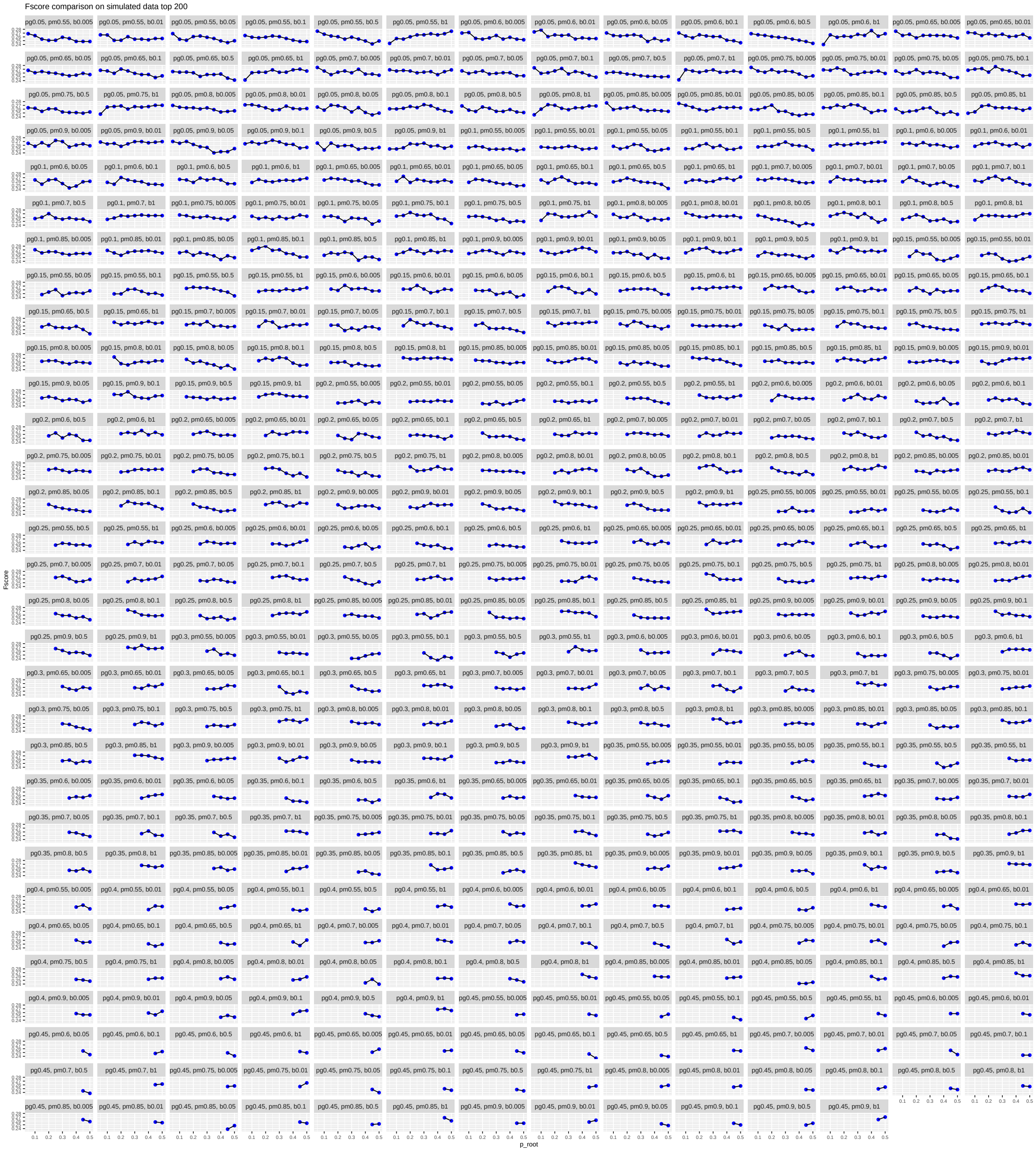

**Supplementary Figure 47.** Effect of $p_r$ parameter on the overall F-score. Changing the $p_r$ parameter while keeping the other parameters fixed, the average performance across 3 cell types based on F-score of top 200 edges on simulated dataset 1.

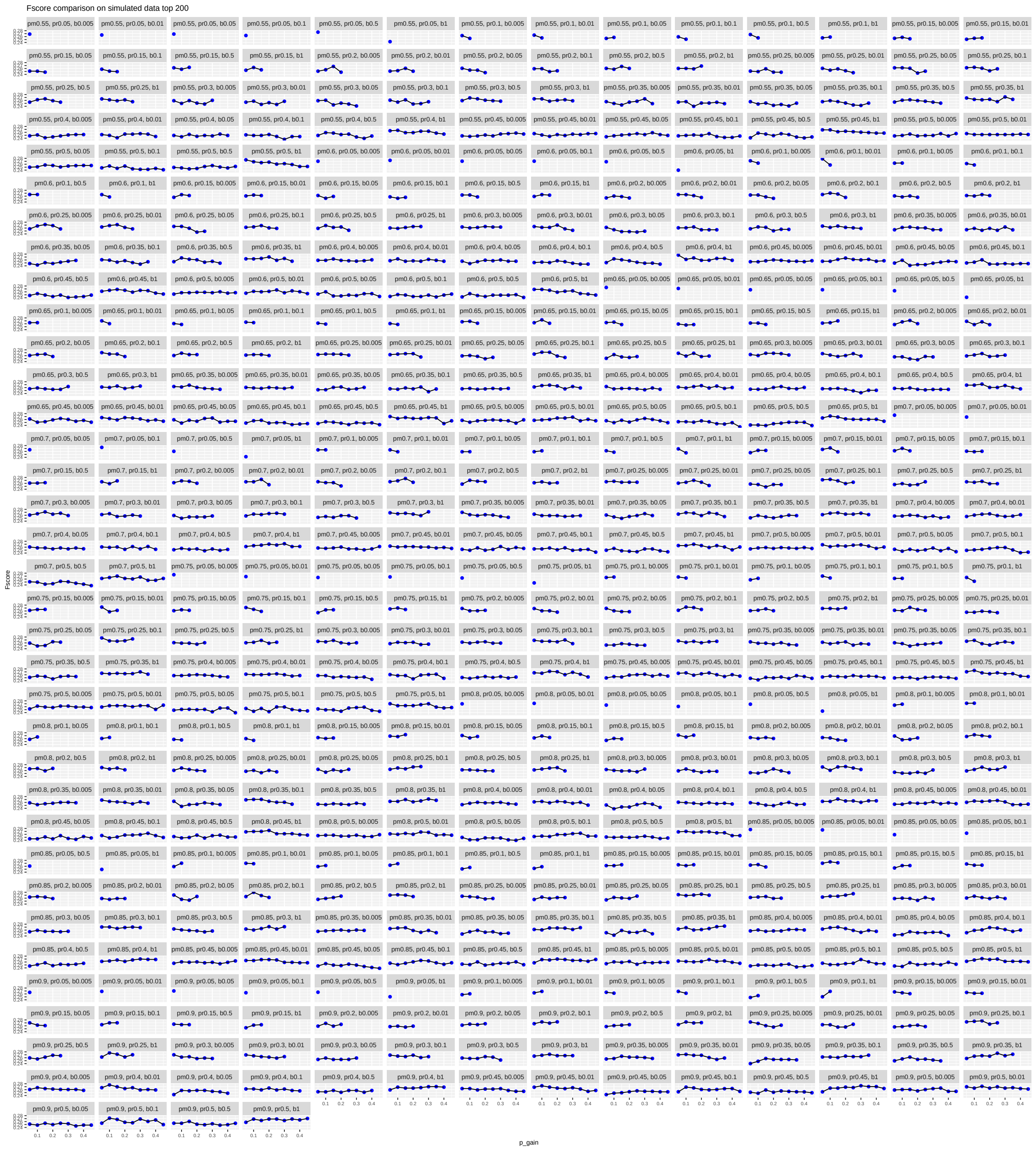

**Supplementary Figure 48.** Effect of $p_g$ parameter on the overall F-score. Changing the $p_g$ parameter while keeping the other parameters fixed, the average performance across 3 cell types based on F-score of top 200 edges on simulated dataset 1.

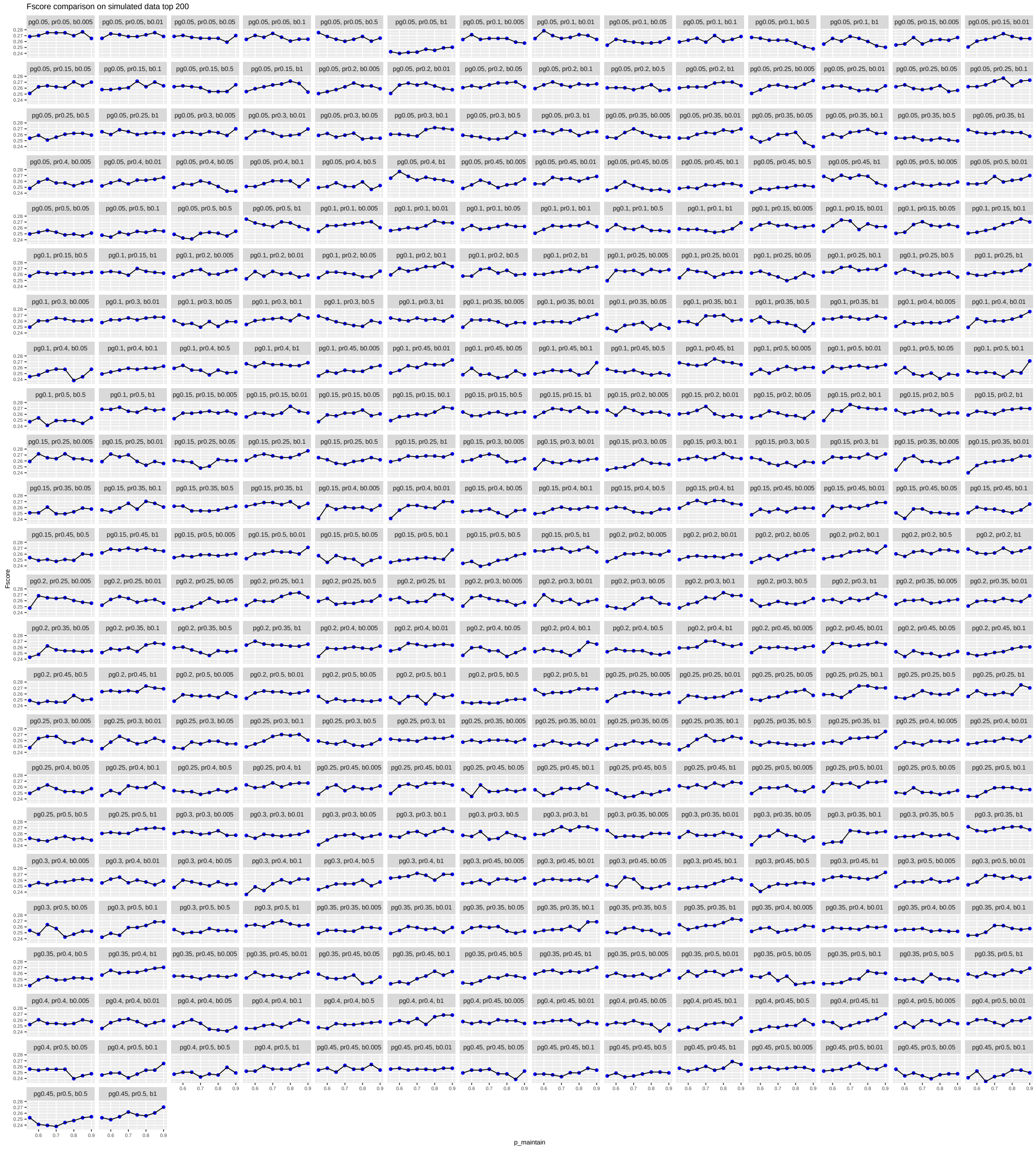

**Supplementary Figure 49.** Effect of $p_m$ parameter on the overall F-score. Changing the $p_m$ parameter while keeping the other parameters fixed, the average performance across 3 cell types based on F-score of top 200 edges on simulated dataset 1.

### Supplementary Methods

#### scMTNI learning algorithm

scMTNI learns the graphs for all cell types simultaneously using a greedy graph learning algorithm with a score-based approach. The score is composed of the data likelihood as well as the structure prior (**Equation 1**).

$$\mathrm{P}(\mathbf{G}, \boldsymbol{\Theta}|\mathbf{D}) \propto \mathrm{P}(\mathbf{D}|\mathbf{G}, \boldsymbol{\Theta})\mathrm{P}(\boldsymbol{\Theta}|\mathbf{G})\mathrm{P}(\mathbf{G}) \tag{1}$$

Our structure learning algorithm begins with $M$ empty graphs for the $M$ cell types and proposes edge additions of possible connections between regulators and targets (**Algorithm 1**). At each iteration, the algorithm scores a candidate edge from regulator $u$ to target gene $v$ based on the change in pseudo likelihood in **Equation** (2) and prior distribution $\mathrm{P}(\mathbf{T})$ in **Equation** (3) and $\mathrm{P}(\mathbf{S})$ in **Equation** (5), and selects the one that corresponds to a local optima. The cell type-specific score improvement of adding an edge from regulator $u$ to target gene $v$ for cell type $d$ is computed as below:

$$\Delta \mathrm{Score}^{(d)}_{u \to v} = \log \mathrm{P}(X^{(d)}_v | \mathbf{R}^{(\mathbf{d})}_{\mathbf{X_v}} \cup X^{(d)}_u) - \log \mathrm{P}(X^{(d)}_v | \mathbf{R}^{(\mathbf{d})}_{\mathbf{X_v}}) + \Delta \log \mathrm{P}(T^{(d)})_{u \to v}. \tag{2}$$

The first two terms are the change in pseudo likelihood, where $\log \mathrm{P}(X^{(d)}_v | \mathbf{R}^{(\mathbf{d})}_{\mathbf{X_v}})$ is the conditional log likelihood of $X^{(d)}_v$ given its neighbor set $\mathbf{R}^{(\mathbf{d})}_{\mathbf{X_v}}$ in cell type $d$'s current graph. The third term is the change of the structure prior, computed as:

$$\Delta \log \mathrm{P}(T^{(d)})_{u \to v} = \log \mathrm{P}(I^{(d)}_{u,v} = 1) - \log \mathrm{P}((I^{(d)}_{u,v} = 0). \tag{3}$$

Since each $I^{(d)}_{u,v}$ can take 1 or 0, there are totally $2^M$ possible combinations of $(I^{(1)}_{u,v}, ..., I^{(M)}_{u,v})$. To make it efficient, we constrain the candidate configurations that meets the following criteria. Based on the cell lineage tree, scMTNI first generates all paths starting from the root node to each leaf node. For each path, it allows up to 1 transition from edge status of 0 to 1 or 1 to 0. For case with 1 transition, we set root cell type status to be $x$, choose one of the intermediate cell types to be $1 - x$, set its descendants to $1 - x$, and set its ancestors to be the same as root. To capture cell type-specific regulation, we also add cell type-specific configurations (i.e. edge present only in one cell type) to the candidate configurations. Among these candidate configurations of edge status across $M$ cell types $(I_{u,v}^{(1)}, ..., I_{u,v}^{(M)})$, we computed the score improvement for each configuration and selected the one that gives the highest score improvement computed as below:

$$\Delta \text{Score}_{u \to v} = \sum_{d=1}^{M} (\Delta \text{Score}_{u \to v}^{(d)} * I_{u,v}^{(d)}) + \Delta \log \text{P}(\mathbf{S})_{u \to v}, \quad (4)$$

where $\Delta \log \text{P}(\mathbf{S})_{u \to v}$ is computed using (5) as the difference of the prior distribution of current configuration and the prior distribution of the previous configuration before adding this edge.

$$\Delta \log \text{P}(\mathbf{S})_{u \to v} = \text{P}(I_{u,v}^{(1)}, ..., I_{u,v}^{(d)} = 1, ..., I_{u,v}^{(M)}) - \text{P}(I_{u,v}^{(1)}, ..., I_{u,v}^{(d)} = 0, ..., I_{u,v}^{(M)}) \quad (5)$$

**Algorithm 1** scMTNI algorithm

**Input**:
- Cell lineage tree $\tau$ with $M$ cell types/clusters
- single-cell expression data $\mathbf{D^{(d)}} = \{\mathbf{X_1^{(d)}}, ..., \mathbf{X_N^{(d)}}\} \in \mathbf{R}^{N \times C}$ for each cell type/cluster $d$
- Structure Prior parameters $\mathbf{S}$: $p_r, p_g^{(d)}, p_m^{(d)}$
- Prior parameters for cell-type specific prior, $\beta_0, \beta_1$
- Prior network for each cell type $\text{P}(\mathbf{T})$

**Output**:
- $M$ cell type-specific gene regulatory networks for each cell type

**Algorithm**:

**while** not converged **do**
    **for** $X_v$ in target gene set **do**
        **for** $X_u$ in regulator set **do**
            **for** cell $d$ in cells **do**
                Compute score improvement of adding this edge $X_u^{(d)} \to X_v^{(d)}$ to the current network in cell type $d$, $\Delta \text{Score}_{u \to v}^{(d)}$
            **end for**
            Compute score improvement of different configurations $\Delta \text{Score}_{u \to v}$ for the edge $X_u^{(d)} \to X_v^{(d)}$ in all cell types, select the configuration $c$ with the highest score.
        **end for**
        Select the regulator $X_{\tilde{u}}$ that gives the highest score improvement and record its configuration $\tilde{c}$.
        **for** cell $d$ in cells **do**
            **if** the edge status for cell type $d$ in configuration $\tilde{c}$ is 1 **then**
                Add edge $X_{\tilde{u}}^{(d)} \to X_v^{(d)}$ to the network of cell type $d$.
            **end if**
        **end for**
    **end for**
**end while**

#### Sensitivity of scMTNI to different parameter settings

We examined how the different parameters affect the overall AUPR and F-score by varying one parameter and keeping the others fixed. For AUPR, decreasing the sparsity parameter $\beta_0$ (i.e. increasing the penalty on adding new edges, $|\beta_0|$ is shown in the figures ) while keeping the other parameters fixed, the average performance across cell types will generally go up slowly for majority of the $(p_g, p_m, p_r)$ settings (**Supplementary Figure 42**). A few exceptions are when $p_g$ and $p_r$ are low, which is likely because the networks being learned are too sparse. However, the overall impact of AUPR was minimal and ranged from 0.25 to 0.26. Increasing the $p_r$ parameter while keeping the other parameters fixed, the average performance generally decreased slightly for settings with smaller $p_g$ and stayed the same for $p_g \geq 0.2$ (**Supplementary Figure 43**). This is also likely due to the fact that the networks are too sparse with a low $p_g$ and low $p_r$. However, here as well the overall change in AUPR was not substantial, ranging from 0.245 to 0.265. Increasing the $p_g$ parameter while keeping the other parameters fixed, the average performance across cell types does not change substantially; it decreases with increasing $p_g$ for some settings (e.g., $\beta_0 = -0.01$,$p_m = 0.6$,$p_r = 0.3$) but remain unchanged for most of the settings (**Supplementary Figure 44**). Finally, increasing the $p_m$ parameter while keeping the other parameters fixed, the average performance across cell types stayed largely unchanged for most of the settings with a few exceptions for higher $p_r$ and $p_m$ where there was a small boost in performance (**Supplementary Figure 45**).

Next, we computed F-score of top 200 edges and examined how the different parameters affect the overall F-score. The F-score helped to more uniformly compare across different settings as the overall density of the inferred edges can affect performance. Based on F-score, decreasing the sparsity parameter $\beta_0(\beta_0 < 0)$ had a less discernable pattern and in most cases the performance remained the same. For some cases, there was increase in performance for higher sparsity, especially when $p_r$ or $p_m$ was high (**Supplementary Figure 46**). Increasing the $p_r$ parameter while keeping the other parameters fixed, the average performance across cell types changed in a manner similar to AUPR with a small downward trend with increasing $p_r$ when $p_g$ was low (**Supplementary Figure 47**). Increasing the $p_g$ parameter affected the average performance in a manner similar to AUPR, with a slight decrease in performance across cell types but remaining unchanged for most of the settings (**Supplementary Figure 48**). Finally, increasing the $p_m$ parameter generally did not affect the average performance for most of the settings with a few exceptions for

| Algorithm | Parameter | Dataset 1 | Dataset 2 | Dataset 3 |
|---|---|---|---|---|
| scMTNI | $p_r$ | 0.1 | 0.2 | 0.4 |
| | $p_g$ | 0.1 | 0.15 | 0.35 |
| | $p_m$ | 0.85 | 0.65 | 0.9 |
| | $\beta_0$ | -0.005 | -0.005 | -0.01 |
| MRTLE | $p_g$ | 0.15 | 0.2 | 0.4 |
| | $p_m$ | 0.6 | 0.55 | 0.85 |
| | $\beta_0$ | -0.01 | -0.05 | -0.01 |
| GNAT | $\lambda_s$ | 35 | 22 | 7 |
| | $\lambda_p$ | 35 | 22 | 5 |
| Ontogenet | $\lambda$ | 2500 | 1000 | 500 |
| | $\gamma$ | 2250 | 1000 | 525 |
| | $\kappa$ | 10 | 1 | 10 |
| INDEP | $\beta_0$ | -0.5 | -0.05 | -0.1 |
| LASSO | $\lambda$ | 0.03 | 0.04 | 0.06 |

**Supplementary Table 1.** Best parameters used for different algorithms on the simulation dataset

higher $p_r$ and $p_m$ where there was a small boost in performance (**Supplementary Figure 49**). In general, the sparsity parameter $\beta_0$ and $p_r$ parameter affected the performance the most, while $p_g$ and $p_m$ parameters did not affect the performance substantially. Overall performance of scMTNI was stable across different parameter configurations (**Supplementary Figure 36**). To compare against methods, we used values across the all different parameter settings (**Supplementary Figure 1, 2**) as well as the best parameter setting (**Supplemental Table 1**, **Supplementary Figure 1, 2**).

| Dataset | # genes | # regulators | # cell types | Avg cells | Memory | | Runtime | |
|---|---|---|---|---|---|---|---|---|
| | | | | | scMTNI | scMTNI+Prior | scMTNI | scMTNI+Prior |
| Cellular reprogramming | 12,216 | 2,036 | 7 | 329 | 1.03±0.50 | 0.90±0.359 | 0.805±0.20 | 0.361±0.106 |
| Adult hematopoiesis | 11,994 | 1,999 | 8 | 346 | 3.06±0.794 | 2.11 ±0.594 | 0.637±0.194 | 0.379±0.80 |
| Fetal hematopoiesis | 16,737 | 2,195 | 11 | 255 | 0.523±0.043 | 0.519±0.138 | 0.632 ±0.325 | 0.286±0.142 |

**Supplementary Table 2.** Characteristics of real datasets and runtimes of scMTNI and scMNTI+prior. Memory (GB) and runtime (hrs) for 50 genes per run for all cell types together in mouse cellular reprogramming data, human hematopoietic data from Buenrostro et al. , and human fetal hematopoiesis data from Ranzoni et al. For memory and runtime, data are presented as mean value +/- SD (number of experiments for cellular reprogramming: $n = 12250$, adult hematopoiesis: $n = 12000$, fetal hematopoiesis:$n = 16750$).
